# TruSight Oncology 500: Enabling Comprehensive Genomic Profiling and Biomarker Reporting with Targeted Sequencing

**DOI:** 10.1101/2020.10.21.349100

**Authors:** Chen Zhao, Tingting Jiang, Jin Hyun Ju, Shile Zhang, Jenhan Tao, Yao Fu, Jenn Lococo, Janel Dockter, Traci Pawlowski, Sven Bilke

## Abstract

**Background:** As knowledge of mechanisms that drive the development of cancer grows, there has been corresponding growth in therapies specific to a mechanism. While these therapies show improvements in patient outcomes, they can be expensive and are effective only for a subset of patients. These treatments drive interest in research focused on the assignment of cancer therapies based on aberrations in individual genes or biomarkers that assess the broader mutational landscape, including microsatellite instability (MSI) and tumor mutational burden (TMB).

**Methods:** Here we describe the TruSight™ Oncology 500 (TSO500; Research Use Only) bioinformatics workflow. This tumor-only approach leverages the next-generation sequencing-based assay TSO500 to enable high fidelity determination of DNA variants across 523 cancer-relevant genes, as well as MSI status and TMB in formalin-fixed paraffin-embedded (FFPE) samples.

**Results:** The TSO500 bioinformatic workflow integrates unique molecular identifier (UMI)-based error correction and a dual approach variant filtering strategy that combines statistical modeling of error rates and database annotations to achieve detection of variants with allele frequency approaching 5% with 99.9998% per base specificity and 99% sensitivity in FFPE samples representing a variety of tumor types. TMB determined using the tumor-only workflow of TSO500 correlated well with tumor-normal (N =170, adjusted *R*^2^=0.9945) and whole-exome sequencing (N=108, adjusted *R*^2^=0.933). Similarly, MSI status determined by TSO500 showed agreement (N=106, 98% agreement) with a MSI-PCR assay.

**Conclusion:** TSO500 is an accurate tumor-only workflow that enables researchers to systematically characterize tumors and identify the next generation of clinical biomarkers.

## Background

It is well established that cancer is a diverse group of diseases resulting from genetic changes that facilitate aberrant and uncontrolled cellular proliferation. Indeed, as the number of known cancer-related genes and understanding of their mode of action grows, there has been a progressive shift towards assigning therapies based on the gene variants present in the tumor of a patient. The growing understanding of the role of genetics in cancer has been enabled by advancements in next-generation sequencing (NGS) technologies that have made the assessment of tumor-related gene alterations easier and more economically feasible. As the number of gene-based therapies and the evidence supporting their increased efficacy grows, there is rising demand for solutions that enable the development of accurate biomarkers that can identify patients most likely to benefit from these therapies in a personalized manner. Biomarkers range from variants in individual genes (eg. BRAF V600E, EGFR) (1–4) as well as others that take into account the broader mutational landscape of a tumor.

Detecting genetic variants in tumors can be difficult due to technical artifacts associated with FFPE samples, which are routinely used in translational research settings. The FFPE process can introduce DNA damage such as deamination (5). Additionally, the amount of tumor content may be a small fraction of the total FFPE sample, and so variants of interest may occur at low frequencies. The detection of somatic variants is further complicated by the high frequency of germline variants (hundreds of mutations per megabase) relative to somatic variants (tens of mutations per megabase) and technical artifacts (over one thousand per megabase) (5–8). While germline variants can be identified by sequencing matched tumor/normal samples, this considerably increases the cost of using NGS for tumor profiling, which remains an important consideration for translational researchers (9).

Microsatellite instability (MSI) status determined using immunohistochemistry (IHC) and polymerase chain reaction (PCR), was the first tissue site-agnostic biomarker approved by the United States Food and Drug Administration for use in selecting patients for immunotherapy. (10,11). Microsatellites are small regions, tens to hundreds of bases in length, of short repeating DNA motifs (1-9 nucleotides). Microsatellite instability is an increase or decrease in the number of these repeats resulting from a mutation in one of the mismatch repair genes - MLH1, MSH2, MSH6 and PMS2 (12,13). Le et al (14,15) were the first to report that mismatch repair-deficient colorectal cancer patients, identified by MSI status, responded better to checkpoint inhibitors. Immunohistochemistry and PCR are established technologies for assessing MSI status (16,17). PCR approaches typically measures the length of five loci, which were first recommended by the Bethesda panel, in the tumor as well as matched normal tissue (16). IHC assays measure the protein level of the mismatch repair genes. Emerging NGS methods for determining MSI status assess many more microsatellite loci than PCR, which improves sensitivity and does not require a matched normal sample (18).

The prevalence of MSI varies between cancers and there are some cancer types that are mostly microsatellite stable, indicating a need for additional biomarkers to complement MSI status (19). Tumor mutational burden (TMB), which reflects the number of tumor mutations per megabase (Mb) of DNA, has been proposed as an additional biomarker. Studies have shown a correlation between higher TMB and the effectiveness of checkpoint inhibitor immunotherapies.(20–22) The current gold standard for determining TMB is whole exome sequencing (WES), but uptake is limited by the cost and long turn-around-time (23). Thus, there is significant interest in the use of targeted panels to determine TMB. There are a number of key factors that influence TMB calculation: panel size and composition, read depth and coverage, assay limit of detection (LoD), and the accuracy of the variant calls (24). Studies have shown that panels over 1 Mb are generally suitable for TMB determination (24,25). With smaller panels, the confidence intervals are too large at lower TMB values (0-30 mutations/Mb) to allow clear distinction between TMB low and TMB high samples. At present there is no established consensus around thresholds for TMB-based therapy selection. Further, the process by which TMB is calculated is not standardized and the bioinformatics behind different commercial assays are a veritable black box.

Here, we describe the bioinformatics workflow of TruSight™ Oncology 500 (TSO500), a pan-cancer NGS assay covering 523 cancer-relevant genes (1.94 Mb) that enables determination of DNA and RNA somatic variants as well as the immuno-oncology relevant biomarkers, MSI and TMB, from a FFPE tumor sample only. The TSO500 bioinformatic workflow leverages unique molecular identifier-based error correction and a statistical model of error rates to suppress technical noise arising from sequencing and FFPE artifacts. To exclude germline variants, the TSO500 workflow uses a database in combination with information about proximal variants. These combined strategies allow the TSO500 workflow to achieve high specificity and sensitivity for detecting variants with low allele frequencies in FFPE samples. Exclusion of technical noise and germline variants also enables TSO500 to determine MSI status and TMB scores that are highly concordant with gold standard approaches.

## Methods

### Sequencing

FFPE samples from a total of 170 patients, and included a range of tumor types: colon, gastric, lung, melanoma, uterine, and endometrial. DNA was extracted using the QiAMP DNA FFPE Tissue Kit (QIAGEN). Manufacturer recommendations were followed for quantitative and qualitative evaluation of DNA quality. Nucleic acids were quantified using a Qubit (Invitrogen, Carlsbad, CA). DNA quality was assessed by a Nanodrop spectrophotometer, with an OD 260/280 value between 1.7 and 2.2 considered acceptable. All samples had at least 40 ng/uL of input DNA.

Library preparation was performed using the hybrid capture-based TruSight Oncology 500 Library Preparation Kit (Illumina, San Diego, CA) following the manufacturer’s protocol. DNA was fragmented to 90 to 250 bp, with a target peak of around 130 bp. Samples then underwent end repair and A-tailing. Next, adapters containing UMIs were ligated to the ends of the DNA fragments. After a purification step, the DNA fragments were amplified using primers to add index sequences for sample multiplexing (required for cluster generation). For TSO500 samples, two hybridization/capture steps were performed, and only one hybridization and capture step was performed for WES samples. First, either a pool of oligos specific to the 523 genes targeted by TSO500 (TSO500 samples) or probes from the IDT xGen Exome panel (WES samples) were hybridized to the prepared DNA libraries overnight. Next, streptavidin magnetic beads (SMBs) were used to capture probes hybridized to the targeted regions. The hybridization and capture steps were repeated using the enriched DNA libraries to ensure high specificity for the captured regions. Primers were used to amplify the enriched libraries before purification using sample purification beads. The enriched libraries were quantified and each library was normalized to ensure a uniform representation in the pooled libraries.

Finally, the libraries were pooled, denatured, and diluted to the appropriate loading concentration. TSO500 libraries were sequenced on a Illumina NextSeq™ 550Dx with a read length of 2×101 bp. Up to 8 TSO500 libraries were sequenced per run. WES libraries were sequenced on an Illumina NovaSeq™ 6000 using 2×101 bp reads using the NovaSeq Xp workflow. Six WES libraries were sequenced per flow cell lane for a total of 24 samples per run (S4 flow cells).

### WES alignment and variant calling

Reads were first mapped to the hg19 genome using the Burrows-Wheeler Aligner (26,27). Next, duplicate reads were marked on the basis of the soft-clip adjusted position and orientation of each read. For a set of duplicate reads, the read with the highest total base quality score was selected as the representative read. Using paired tumor and normal samples, somatic and germline variants were called using Strelka2 with the “--exome” option enabled and default values for all other parameters (28). Both germline and somatic variants were then annotated using Nirvana (29).

### TSO500 alignment and variant calling

A diagram of the TSO500 bioinformatics pipeline is provided in **Figure 1**; quality control metrics are shown in **Supplemental Table 2**. Data analysis begins with an initial DNA alignment to the hg19 genome using the Burrows-Wheeler Aligner. Next using non-random unique molecular identifiers (NRUMIs), duplicate reads are collapsed to reduce PCR and sequencing errors. In the meantime, read collapsing increases the base quality of real low frequency variations, leading to high sensitivity to call low frequency mutations. The NRUMIs used in TSO500 were designed with a pair-wise edit distance of at least 3, which allows error correction of the UMI itself with up to 2 mismatches/indels. Due to A-tailing during library preparation, there would be a lack of nucleotide heterogeneity at the attachment point if a single length UMI was used, so the method uses variable length UMIs (6 and 7-mers) to increase nucleotide heterogeneity: the 7^th^ base is T for all 6-mers, and for the 7-mers the 7^th^ base is never T. NRUMIs are designed to have a 3bp edit distance between any pair, and so any 1bp error within a NRUMI sequence can be corrected. Families of duplicate reads are identified on the basis of both their aligned position and having matched NRUMIs, which are added to both ends of DNA molecules during sequencing library preparation. Duplicate read families are then collapsed into a single consensus sequence. Collapsed sequences supported by reads from both the forward and reverse strand are denoted as duplex sequences, whereas sequences supported by reads from just one strand are denoted as simplex sequences.

**Figure 1.**
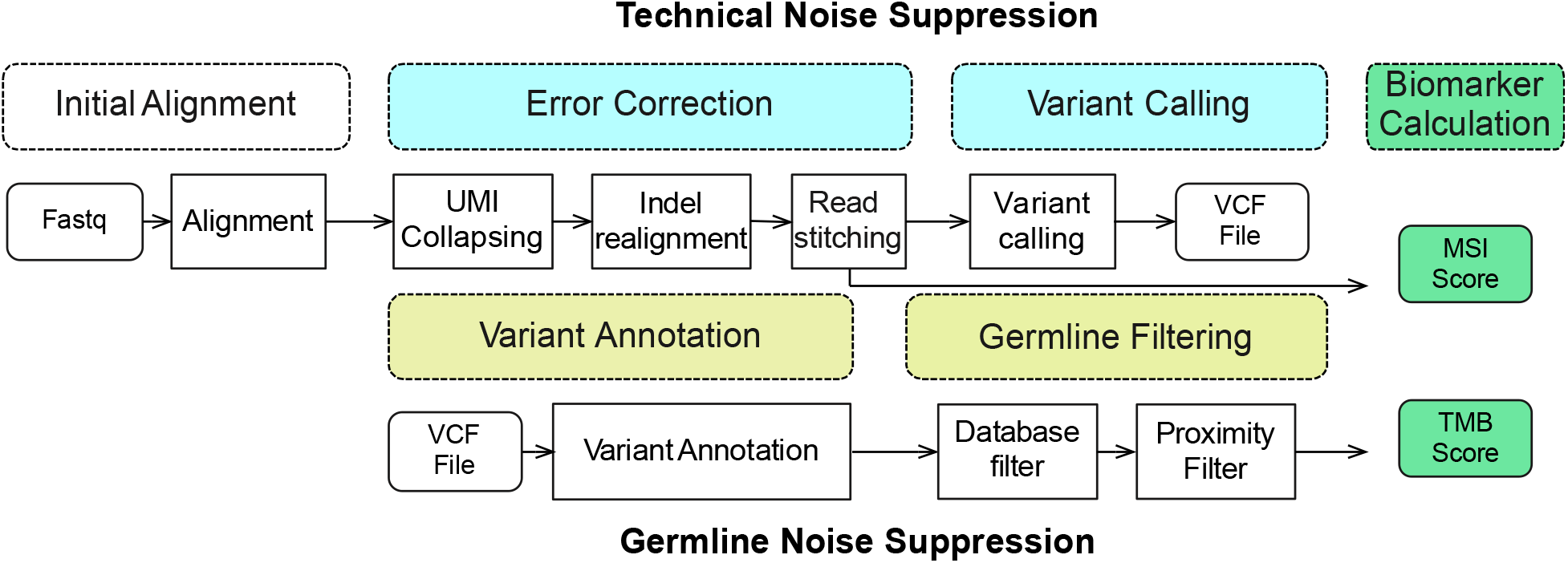
TSO500 bioinformatic workflow for variant calling and biomarker calculation . Abbreviations: Indel = insertion/deletion, MSI = microsatellite instability, TMB = tumor mutational burden, UMI = unique molecular identifier.

NRUMI collapsed sequences are then indel realigned and stitched together into a consensus fragment using Gemini (30). Stitching further improves per base accuracy and quality and allows for a more accurate calculation of coverage in the overlapping region between read pairs. Reads ending near detected indels are realigned to remove alignment artifacts. Somatic variant calling is then performed using Pisces (31) to identify candidate variants. Variants that do not overlap a region targeted by the TSO500 assay are removed. Post-processing of the small variant calls removes variants that have error rates below quality thresholds (described in the following section). The final set of variants is then annotated using Nirvana (29).

### Technical noise suppression

To achieve high fidelity in variant calls, a base-change specific variant filtering algorithm is applied to remove recurrent artifacts and FFPE deamination artifacts. For each variant of interest, background noise at the same site is estimated from ~60 normal FFPE samples profiled by TSO500 assay. Using a binomial distribution to model the background noise level, we then test the significance of each observed mutant allele depth given the total depth and convert the p-value to a variant quality score (AQ) as −10*log_10_ (*p*-value). In addition, the error rate of each nucleotide change is estimated for each support category – duplex sequences (sequences supported by both the forward and reverse strand), simplex sequences originating from forward strand, and simplex sequences originating from reverse strand - in each sample by using all the positions with an allele frequency less than 5%. **Supplemental Figure 1** shows that the error rate can vary considerably depending on support category and nucleotide change. The error rate is calculated as the total mutant reads divided by total depth. For variants of interest, a likelihood ratio score (LQ) is calculated as follows:

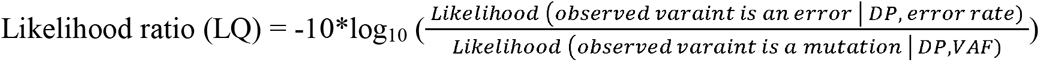

For variants with a Catalogue of Somatic Mutations in Cancer (COSMIC) (32) count > 50, the LQ and AQ thresholds are 20 and the remaining sites are 60.

### Germline variant filtering

To calculate TMB, germline variants are removed through a strategy that leverages information from public databases combined with the coverage and allele frequency of variants in the same region. The database filter uses information about observed variation from multiple population groups to filter common germline variants. Variants that have an observed allele count of N (where the pipeline default is 10) or more in either the 1000 genome, gnomAD exome, or gnomAD genome database, are filtered out (**Supplemental Figure 2**) (33,34).

Database strategies alone are not sufficient for filtering out private germline mutations in each individual. For variants remaining after database filtering, we apply an additional filter, which uses the allele frequency of proximal known germline variants to determine whether a variant is likely a germline variant. For a given variant, we identify variants on the same chromosome with an allele frequency within the greater of 2 standard deviations (assuming a binomial distribution using coverage and allele frequency of the given variant) and 0.05. If there are more than 5 variants total and more than 95% of them are found in the germline database, then the variant of interest is considered to be likely germline. Additionally, variants with an allele frequency ≥ 90% are labeled as likely germline. In **Supplemental Figure 2**, we show the proximal variants (circled variants) that are within allele frequency 0.05 (black bars) for a germline variant not filtered using database allele counts. This is a simple yet effective approach that accounts for both unfiltered variants in expected germline frequency ranges and germline variants with allele frequency shifts due to copy number variant (CNV) events. One caveat of proximity filter is that it may filter out real somatic mutations when the tumor purity is extremely high (>=80%), leading to under-estimation of tumor mutational burden (TMB). We also note that germline filtering is for the purpose of TMB estimation. The classification of germline versus somatic is not perfect, and it is not meant for identification of somatic mutations per se.

### Tumor mutational burden

TMB metrics are calculated following small variant calling. TMB is the ratio of the number of eligible variants (mutations) to eligible DNA regions (Mb). Eligible variants (numerator) include only coding variants with a frequency of ≥ 5% and coverage ≥ 50 reads. Single-nucleotide variants (SNVs) and indels are included, but multi-nucleotide variants (MNVs) and variants with a COSMIC count of ≥ 50 are excluded. Variants in blacklisted regions with poor mappability are also excluded. For the denominator, all eligible coding regions (with coverage ≥ 50x) are included, except for the blacklist regions. The TSO500 bioinformatics workflow outputs both the TMB score and details of the variants that are included in the TMB calculation.

### Microsatellite instability status

TSO500 determines microsatellite instability status from stitched reads from a tumor sample only. Unstable microsatellite sites are detected by assessing the shift in the length of a microsatellite site for a tumor sample against a set of normal baseline samples. First, the baseline Jensen-Shannon distance, *d*, is calculated for each pair of baseline samples *i*, *j* yielding *D*_*i,j*_. Next, Jensen-Shannon distance is calculated between the tumor sample *x* and each baseline sample, yielding *D*_*x,i*_. These two distributions are then compared using a one-sided t-test under the null hypothesis *D*_*i,j*_ ≥ *D*_*x,i*_. Sites with *p* < 0.01 and *D*_*x,i*_ − *D*_*i,j*_ > 0.01 are reported as unstable. We require that each MSI site assessed have at least 60 full-spanning reads. The proportion of unstable MSI sites to total assessed MSI sites is reported as a sample-level microsatellite score, 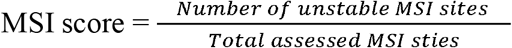. We require at least 40 sites to be assessed to determine a MSI score.

To ensure that microsatellite instability can be assessed in a wide range of tumor types, the TSO500 assay initially targeted 175 noncoding homopolymer regions. We observed that certain homopolymer regions tended to test as unstable more frequently for samples with a particular ethnic background. Using all sites, we observe that African samples tended to have considerably higher MSI scores in comparison to samples from other ethnic backgrounds (**Supplemental Figure 4**, n=101 European, n=23 East Asian, n=10 African, and m=6 Hispanic). After exclusion of sites that showed an ethnicity bias, we observe more comparable MSI scores for samples with different ethnic backgrounds. To ensure robust MSI status calculation, we also exclude sites with low sequencing coverage. Using these two criteria for site selection, we assess a total of 130 noncoding homopolymer regions to determine MSI status.

## Results

### Technical sequencing noise suppression

High accuracy biomarker calculation is dependent on high-accurate sequencing, alignment and variant calling. The TSO500 assay integrates UMIs for error correction, allowing for suppression of PCR duplicates and sequencing errors, while preserving DNA molecules supporting low frequency variants. **Figure 2** presents consolidated error rates before and after UMI read collapsing for 170 normal FFPE samples. We observe a systematic reduction in error rates across all samples that is independent of overall sample error rate. The mean error rate for all bases in the panel was 0.074% before condensing and 0.032% after condensing. UMI based error correction performance improved with greater family depth (**Supplemental Figure 3**). Mean family depth was 4.788 across all samples.

**Figure 2.**
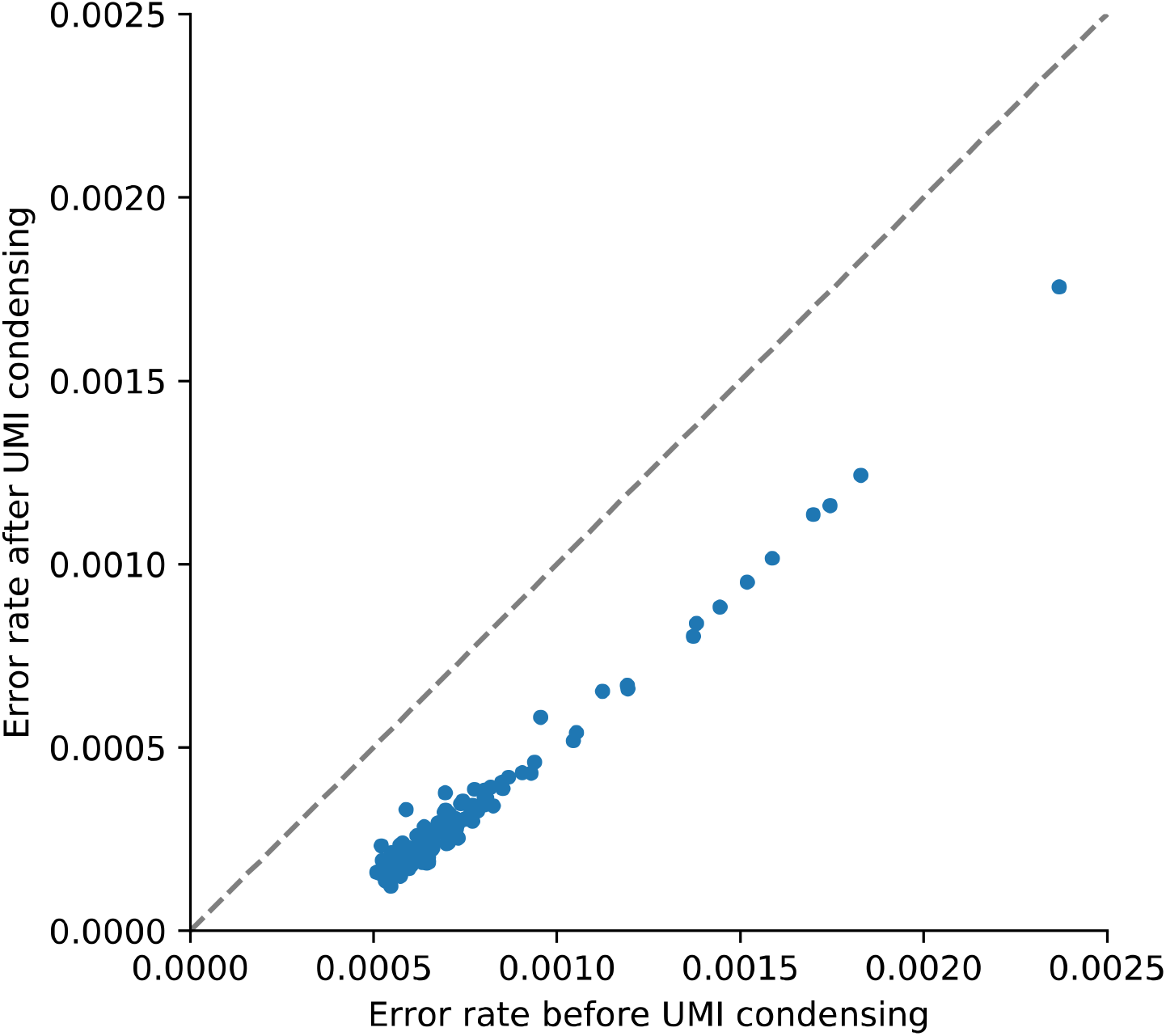
Reduction in error rates using unique molecular identifiers. Sequencing error rates for 170 normal FFPE samples before and after condensing reads using unique molecular identifiers.

UMI based read collapsing suppresses sequencing error, but there are additional sources of false positive calls that are quality-dependent and/or caused by DNA damage. For example, deamination (C:G→T:A) and oxidation (C:G→A:T) events were observed with varying rates depending on the strand and whether a read was simplex or duplex (**Supplemental Figure 1**). Therefore, we use normal FFPE samples to estimate the background noise at each base and model the confidence of each variant call. We also calculate the error rate of each nucleotide change for each sample to estimate a sample-specific error profile (see methods). Using the sample-specific error profile, we calculate a sample-specific variant calling threshold, allowing us to achieve comparable variant calling performance between samples of varying quality. By comparing the likelihood that a variant occurs at a given position against the likelihood that an error has occurred at a given position, we can further eliminate low quality, false positive variant calls. **Figure 3** gives the distribution of total false positive variant calls for 170 normal FFPE samples across the entire 1.94 Mb TSO500 panel; overall, we achieve specificity > 99.9998% in these samples.

**Figure 3.**
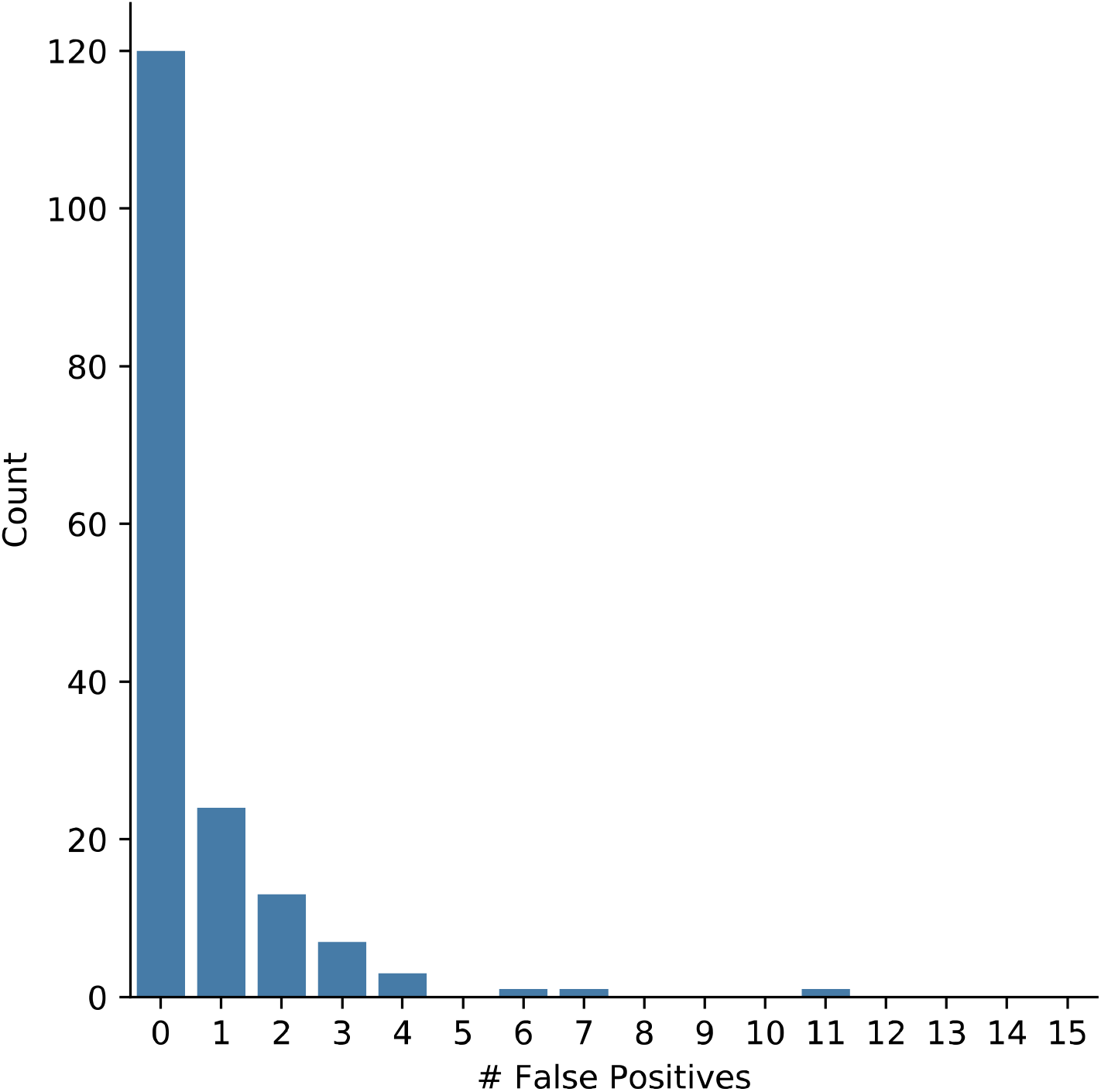
Distribution of the number of putative false positive variant calls in 170 normal FFPE tissue samples. The vertical axis gives the number of samples.

To estimate our limit of detection and sensitivity for small variant calling, we performed a series of dilution experiments using 10 FFPE tissue samples and 4 commercial reference standards. Variant allele frequencies (VAFs) for 35 variants were determined in undiluted samples using digital droplet PCR and ranged from 21-78%. These samples were then diluted so that VAFs ranged as low as 1-2% and that the majority of variants had VAF ≤ 6%. Summary statistics for our variant calling performance in these diluted samples are shown in **Table 1**. For variants with VAF of 5±2.5% we achieve sensitivity of 99%.

**Table 1.** Summary statistics for limit of detection for small DNA variants. Abbreviations: MNV = multi-nucleotide variant, SNV = single-nucleotide variant, VAF = variant allele frequency.

| Expected VAF* | Variant Type | Called Samples / Total Samples | Sensitivity (%) |
| --- | --- | --- | --- |
| 10.0% ± 2.5% | SNV | 18/18 | 100.0 |
|  | Insertion | 4/4 | 100.0 |
|  | Deletion | 4/4 | 100.0 |
|  | Complex Indel | 10/10 | 100.0 |
|  | **All** | **36/36** | **100.0** |
| 5.0% ± 2.5% | SNV | 433/434 | 99.8 |
|  | MNV | 4/4 | 100.0 |
|  | Insertion | 25/26 | 96.2 |
|  | Deletion | 104/108 | 96.3 |
|  | Complex Indel | 10/10 | 100.0 |
|  | **All** | **576/582** | **99.0** |

### Germline variant filtering

As TSO500 is a tumor only workflow, our variant calling algorithms cannot exclude germline variants by using information from a matched normal sample. Instead, identification of germline variants is accomplished by using information from public databases of germline variants (33,34) and a second line proximity filter. By excluding variants that appear more than 10 times in public databases, we were able to identify, on average, 99.73% of the coding region variants as germline variants in each sample; germline variants were verified by assaying the matched normal samples using with TSO500. (33,34). While database filtering is effective, a median of 4 germline variants, not represented in databases, remained unfiltered in our set of 170 tumor samples. And so, we applied an additional filter that uses the database information of proximal variants (see methods). After applying the proximity filter, the number of remaining germline variants in each sample, germline residuals, was further reduced to a median of 1 per sample (**Figure 4**). We calculated and compared TMB scores for each sample using the tumor sample only and using both the tumor and normal sample. The high correlation of the tumor-only TMB values with the matched tumor-normal values (adjusted *R*^2^ = 0.9945) further demonstrates the effectiveness of our germline filters and highlights that our tumor only workflow can accurately estimate TMB (**Figure 5A**).

**Figure 4.**
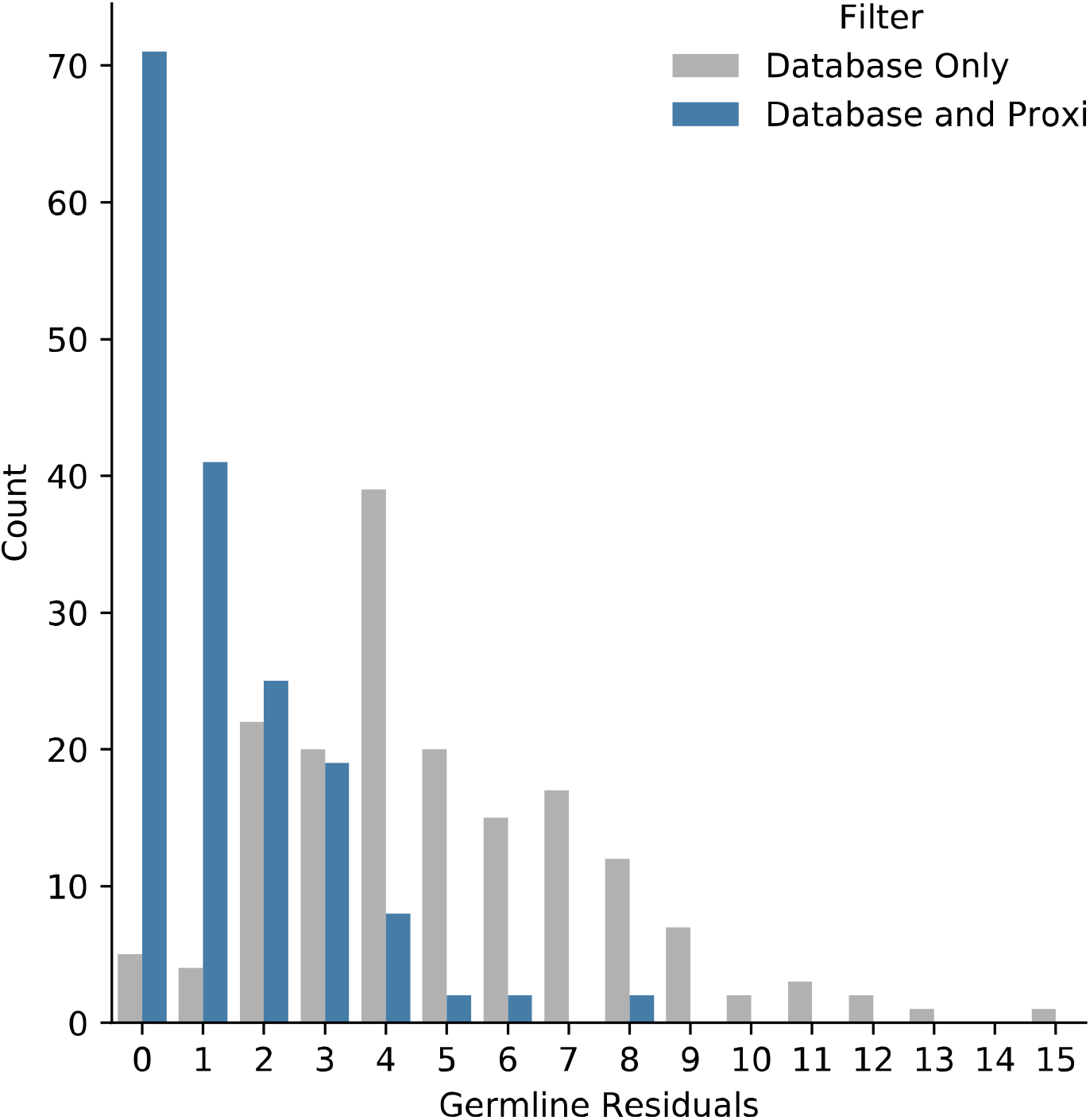
Reduction in putative germline variants using the proximity filter for 170 tumor FFPE tissue samples. Germline variants were determined by comparing to a matched normal FFPE tissue sample. The horizontal axis gives the number germline variants remaining either after database filtering only or after database and proximity filtering. The vertical axis gives the number of samples.

**Figure 5.**
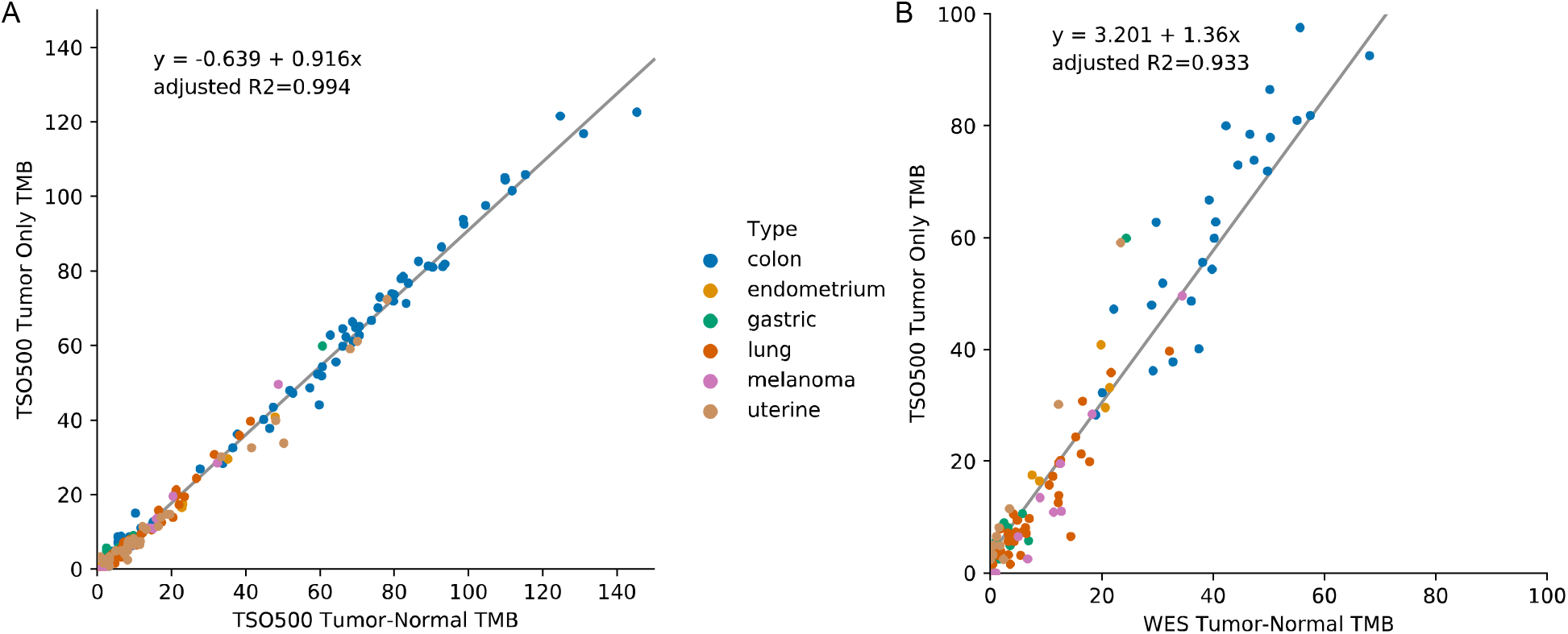
Comparison of TSO500 tumor-only TMB values to tumor-normal TMB values. A. TMB values using the TSO500 workflow are shown for 170 Tumor-Normal pairs. Tumor-only TMB is strongly correlated with Tumor-Normal TMB in formalin-fixed, paraffin-embedded tissue samples (adjusted *r*^2^ = 0.9945). B. TMB values are shown for 108 Tumor-Normal pairs. Tumor-only TSO500 TMB is strongly correlated with WES TMB in formalin-fixed, paraffin-embedded tissue samples (adjusted *r*^2^ = 0.933). TSO500 TMB is calculated using both SNVs and Indels. WES TMB is calculated using nonsynonymous variants only. Abbreviations: TMB = tumor mutational burden, WES = whole exome sequencing.

### Tumor mutational burden using tumor sample only

Given the formulation of TMB, high quality alignment and variant calling with TSO500 should enable robust TMB calculation. The net effect of UMI based read collapsing and technical noise and germline variant filtering is a reduction in false positive variants in a typical FFPE sample from approximately 1500/Mb to < 5/Mb. To assess whether this reduction in false positives translated into reliable TMB values concordant with WES sequencing, we re-processed 108 out of 170 of our samples with whole exome sequencing (matched tumor-normal samples) and calculated the nonsynonymous TMB, which is the typically reported TMB value for WES data (35–37). We then compared the WES tumor-normal TMB values with TSO500 tumor-only TMB values (by default TSO500 uses both coding SNVs and Indels regardless of functional impact, total TMB), observing a high correlation between the two sets of TMB values (adjusted *R*^2^=0.933, **Figure 5B**). We observe that TSO500 TMB values tend to be elevated relative to WES TMB (intercept=3.201), which is likely attributable to the use of total TMB instead of only nonsynonymous variants for WES. Similarly, the elevated slope of the best fit line (1.36) is attributable to the same reason.

We next assessed the reproducibility of TSO500 tumor-only TMB scores. Four FFPE samples and four commercially available control samples were selected to be assayed across a variety of conditions. Using each of these eight FFPE samples, three operators generated libraries using three lots of reagents, three sets of instruments with each combination of reagents in duplicate, totaling to 36 technical replicates per sample. Across all eight samples, we observed highly similar values across all operators and conditions (**Supplemental Figure 6**); the mean coefficient of variation across all samples was 0.0567.

### Microsatellite instability status using tumor sample only

The TSO500 bioinformatics workflow leverages UMI error-corrected stitched reads to determine MSI status from a tumor sample only. To ensure that TSO500 can determine MSI status in a wide range of tumor sites, a total of 130 noncoding homopolymer regions are assessed for microsatellite instability. We also specifically exclude sites demonstrating an ethnicity bias (**Supplemental Figure 4**). To enable a tumor-only determination of MSI status, TSO500 uses an information theory-based approach that compares the distribution of site differences between the tumor sample and a panel of normal samples with the distribution of differences between samples within the panel of normal (see methods). From our initial set of 170 FFPE samples, we assayed 106 samples for MSI status using an MSI-PCR assay commercially available from Promega that assess five mononucleotide sites (**Figure 6**). Using a MSI score cutoff to separate MSI-high from MSI-stable samples, the TSO500 MSI score showed 98% overall agreement with MSI-PCR (**Supplemental Table 3**).

**Figure 6.**
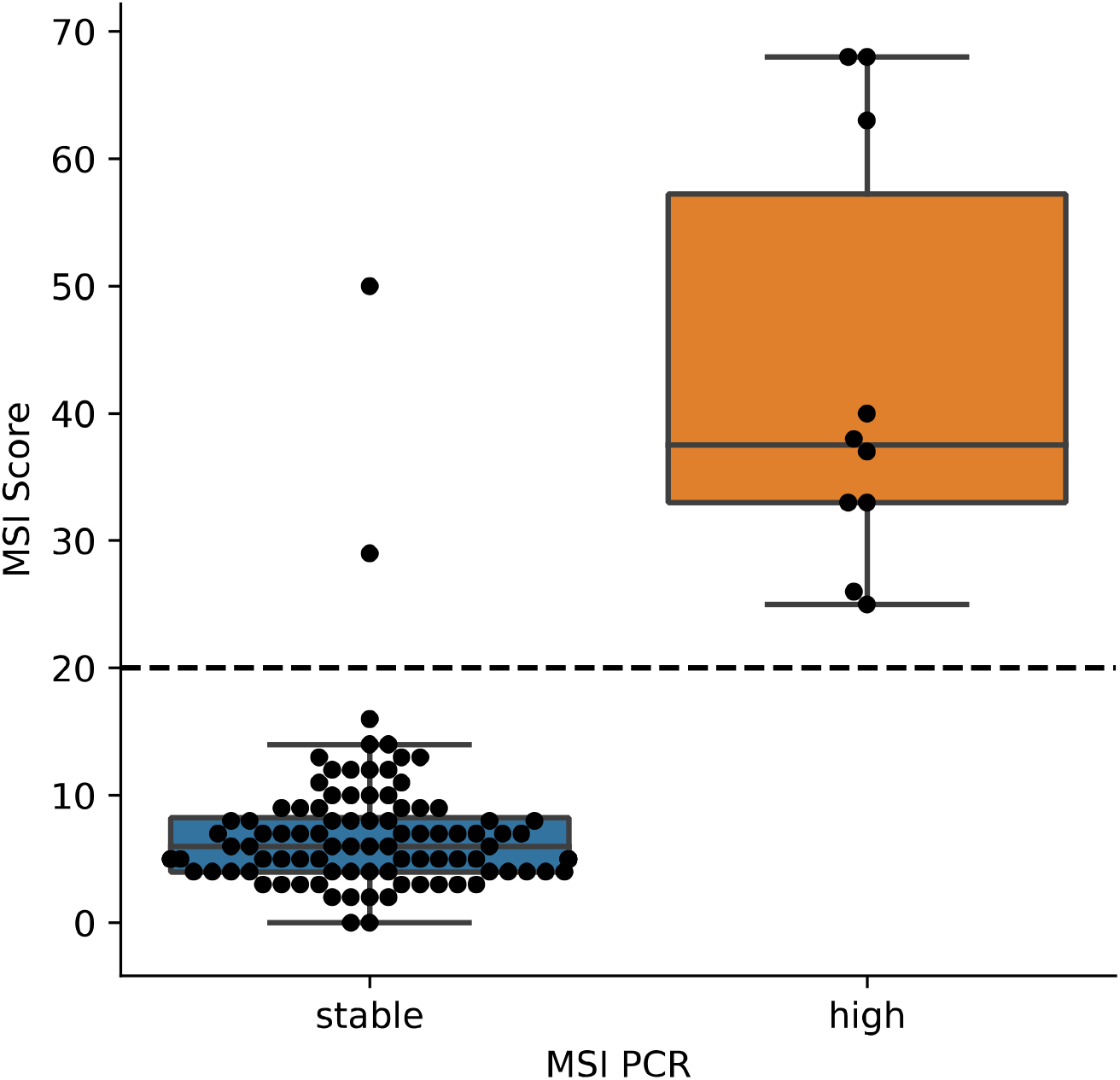
Comparison of tumor-normal MSI-PCR and tumor-only TSO500 MSI-score. MSI status determined by Promega MSI-PCR using matched tumor and normal samples is shown on the horizontal axis for 106 FFPE samples from a variety of tumor types (colon, gastric, lung, melanoma, uterine, endometrium). The vertical axis gives the TSO500 tumor-only MSI score. The dashed line indicates the MSI score cutoff for determining MSI-high samples.

To examine the reproducibility of determining MSI status using the tumor only workflow of TSO500, we performed a total of 36 technical replicate experiments for eight samples (four FFPE samples and four commercially available controls). Each sample was processed by three operators who used three lots of reagents and three sets of instruments to perform each experiment twice. We observed robust MSI scores across all operators and conditions (**Supplemental Figure 7**) and the mean standard deviation across all samples was 0.0196.

## Discussion

The treatment of cancer is moving towards the assignment of treatment based on the genetic attributes of an individual patient’s tumor. Given the wide landscape of cancerous genetic aberrations and treatments specifically targeting a specific mechanism, there is a pressing need for cost effective methods to systematically characterize a tumor. Here we present the bioinformatics solutions of TSO500, a cost-effective tumor only pan-cancer NGS assay, which enables the identification of a range of DNA variant types (including SNVs, MNVs, indels, and complex indels), as well as the determination of MSI and TMB. We describe the bioinformatic algorithms underlying our error correction, variant calling, and determination of biomarkers using a tumor sample only. In addition to showing that tumor only analysis with TSO500 enables accurate tumor variant calls, we also demonstrate that biomarker metrics reported by TSO500 are highly concordant with gold standard methods and are highly reproducible. Our bioinformatics solutions are available as Linux containers (Docker and Singularity images, support.illumina.com/sequencing/sequencing_kits/trusight-oncology-500/questions.html), which enables on-premise software deployment when security is a high priority or when bioinformatics expertise is not readily available (38).

The tumor-only workflow of TSO500 is designed to be effective for research applications today, as well as for the development of future applications. TSO500 enables research into emerging biomarkers such as MSI and TMB status. TMB has been demonstrated to be useful for determining the success of checkpoint inhibitor immunotherapies in NSCLC patients (20–22). Similarly, there is evidence that MSI status may identify CRC patients that respond better to checkpoint inhibitor immunotherapies (14,15,39). Despite the promise of these biomarkers, we note that the National Comprehensive Cancer Network (NCCN) guidelines for using these biomarkers are still emerging, partly because methods for determining these biomarkers are still emerging (40). Here, we fully describe the algorithms underlying our TMB and MSI calculations to enable further research and development of these biomarkers. Furthermore, data from the TSO500 workflow represents a systematic interrogation of the mutational landscape of a tumor and can be leveraged to develop future biomarkers. We are in the process of developing an in vitro diagnostic (IVD) based on TSO500 content. This IVD is anticipated to have the same technical capabilities described here and will enable clinical application of this technology.

We envision several additional applications using TSO500. Several variant types not described in detail here – including RNA variants, copy number variation, and splice variants - can also be detected using TSO500. These additional variant types may contribute to future biomarkers as research progresses (41–44). We are also developing a circulating tumor DNA (ctDNA) version of TSO500 because there is increasing evidence that supports the use of liquid biopsies, and specifically the analysis of ctDNA, to diagnose and monitor the treatment of cancer patients (45–50). Next-generation sequencing enables characterization of each patient’s tumor to a level not previously achievable, and we expect that these technologies will accelerate progress towards the implementation of precision medicine in oncology.

## Conclusion

TSO500 is an accurate tumor-only workflow that enables researchers to systematically characterize tumors and identify the next generation of clinical biomarkers. We envision the broader adoption of comprehensive genomic profiling panels like TSO500 will further help advance precision medicine.

## List of abbreviations

MSI: microsatellite instability
TMB: tumor mutational burden
TSO500: TruSight™ Oncology 500
FFPE: formalin-fixed paraffin-embedded
UMI: unique molecular identifier
NGS: next-generation sequencing
IHC: immunohistochemistry
PCR: polymerase chain reaction
WES: whole exome sequencing
NRUMIs: non-random unique molecular identifiers
AQ: variant quality score
LQ: likelihood ratio score
COSMIC: Catalogue of Somatic Mutations in Cancer
CNV: copy number variant
VAFs: Variant allele frequencies

## Declarations

### Ethics approval and consent to participate

The FFPE samples used in this study were purchased from commercial vendors.

### Consent to publish

Not applicable

### Availability of data and materials

The dataset will be available after being authorized by the corresponding author on reasonable request.

### Competing interests

All of the authors are employees of and own stock in Illumina, Inc.

### Funding

This research was supported by Illumina, Inc.

### Authors’ Contributions

Study concept and design: CZ, JL, JD, TP, SB. Algorithm development and data analysis: CZ, TJ, JHJ, SZ, JT, YF. The authors have read and approved the final manuscript.

## Acknowledgements

None

---

**Supplemental Table 1.**
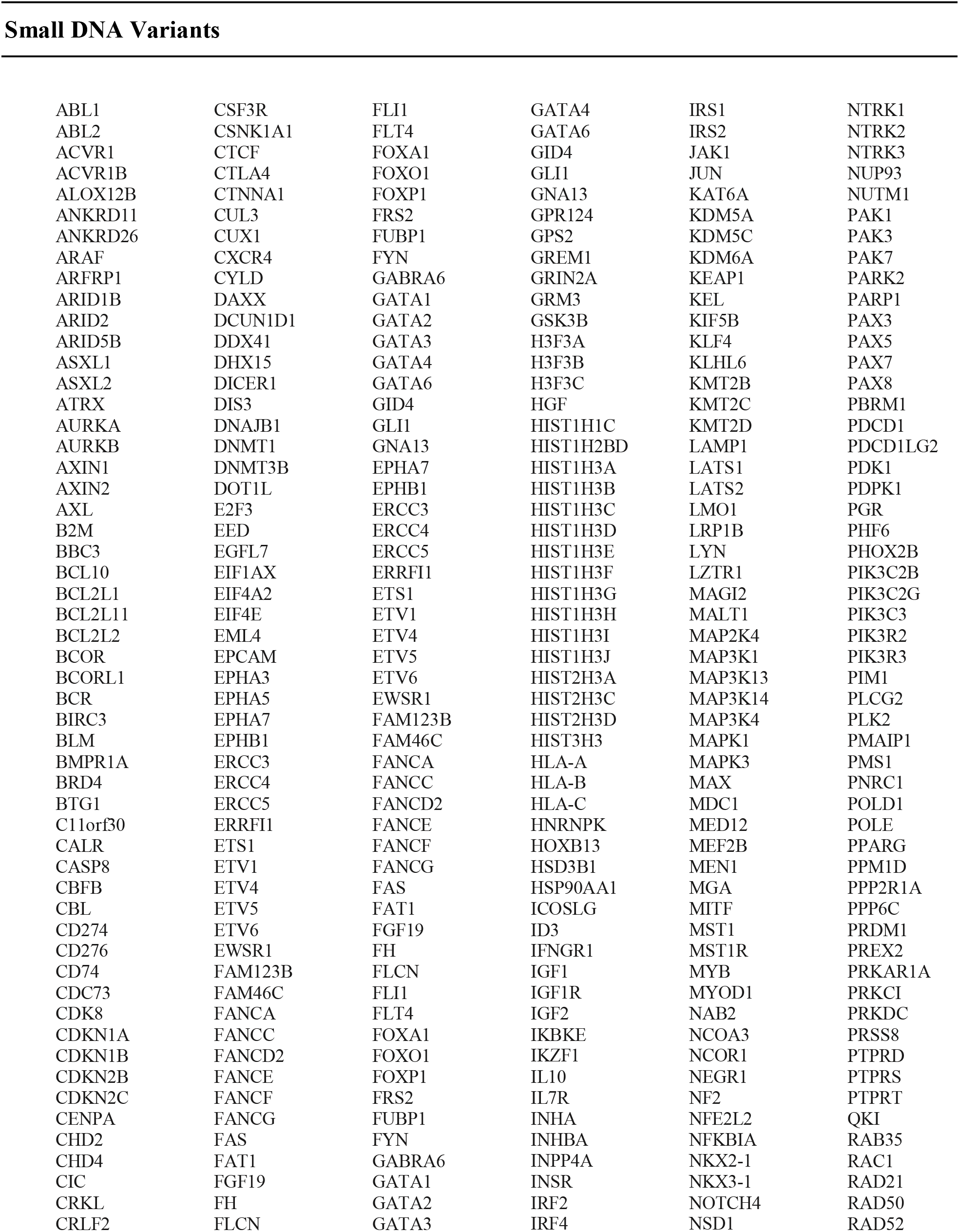
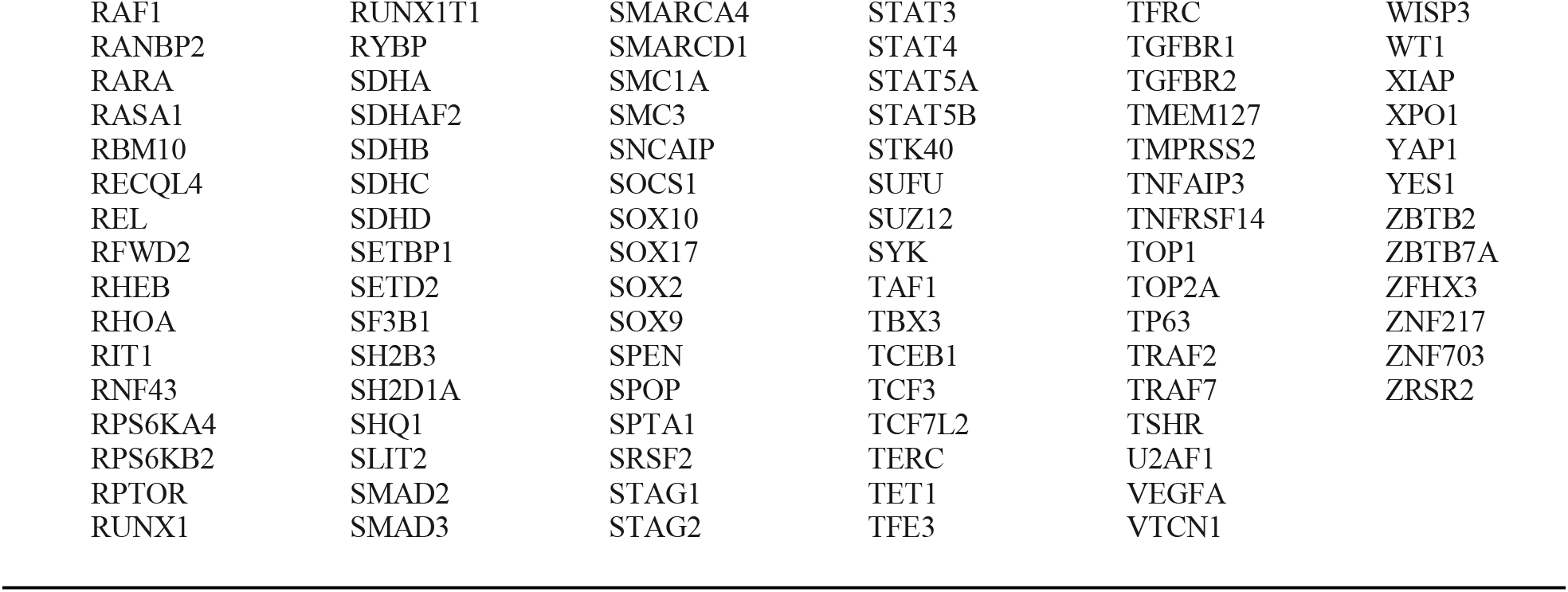
TSO500 assay content (523 genes).

**Supplemental Table 2.**
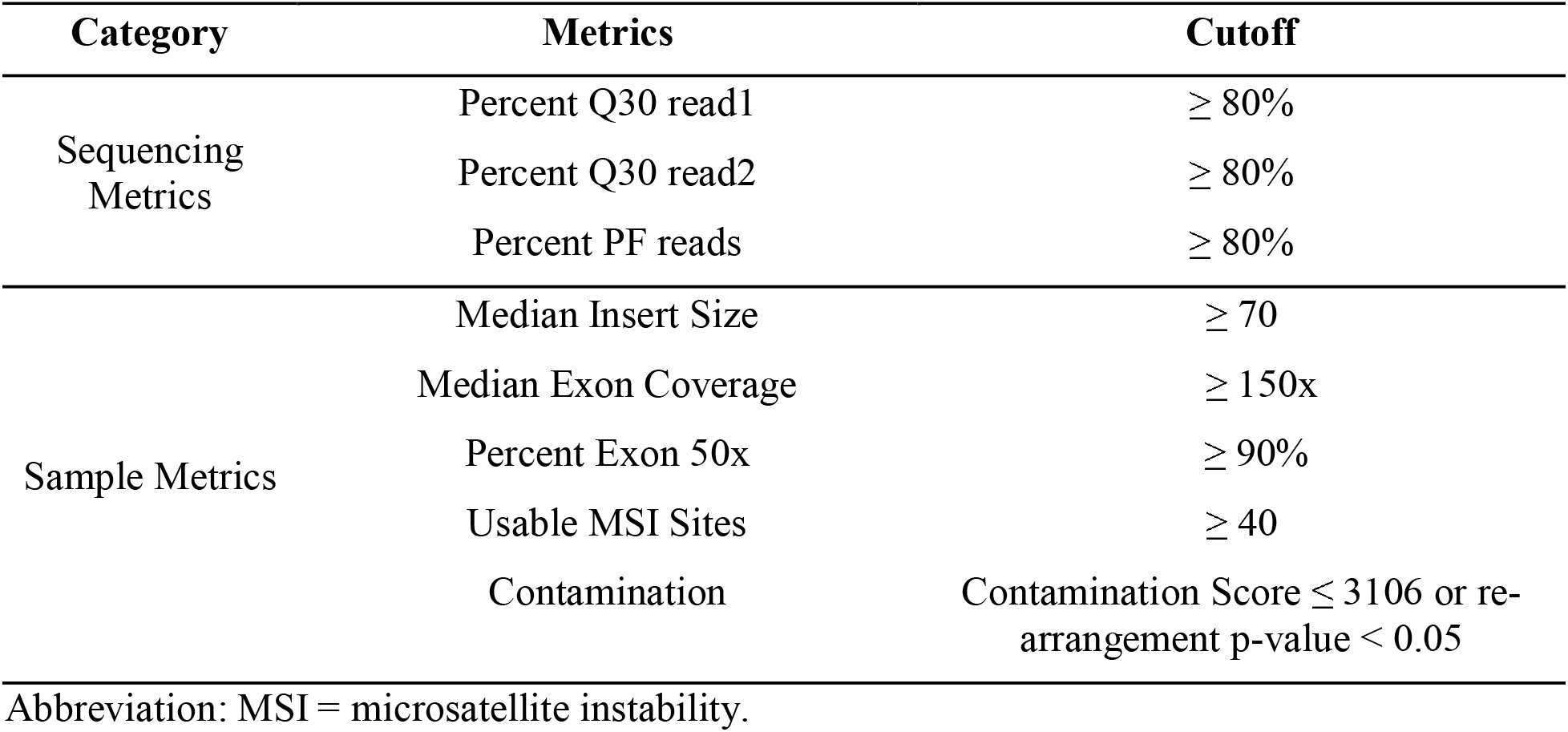
TSO500 quality control metrics.

| Category | Metrics | Cutoff |
| --- | --- | --- |
| Sequencing Metrics | Percent Q30 read1 | $\geq 80\%$ |
| | Percent Q30 read2 | $\geq 80\%$ |
| | Percent PF reads | $\geq 80\%$ |
| Sample Metrics | Median Insert Size | $\geq 70$ |
| | Median Exon Coverage | $\geq 150x$ |
| | Percent Exon 50x | $\geq 90\%$ |
| | Usable MSI Sites | $\geq 40$ |
| | Contamination | Contamination Score $\leq 3106$ or re-arrangement p-value $< 0.05$ |

Abbreviation: MSI = microsatellite instability.

**Supplemental Table 3.** Details of MSI samples.

**Supplemental Figure 1.**
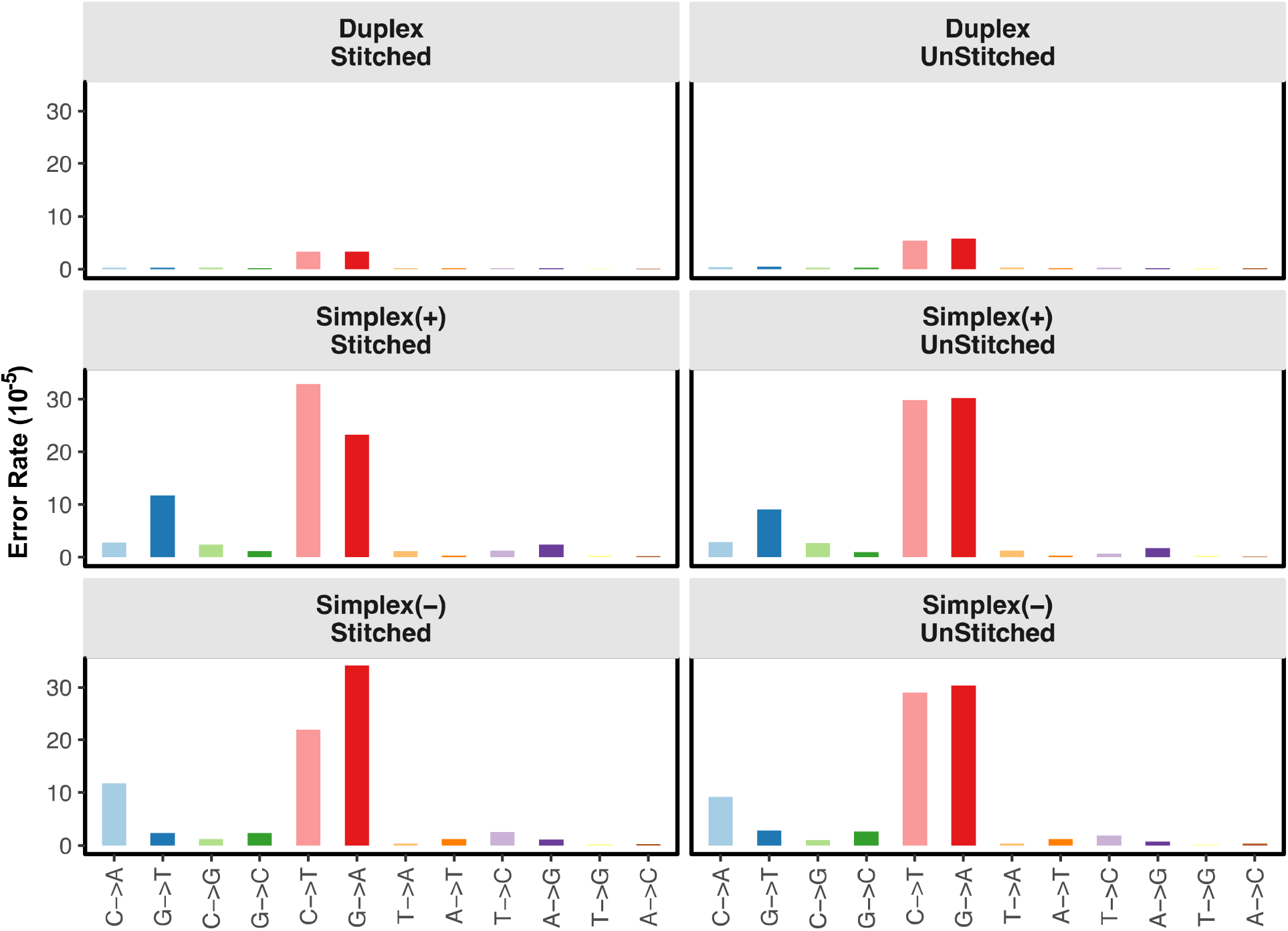
Error rates from the simplex forward, simplex reverse, and duplex strands with and without stitching. Deamination errors (G→A), which are prevalent in formalin-fixed, paraffin-embedded tissue and oxidation errors resulting from with sonication (C→A or G→T errors) are greatly reduced in the duplex, stitched panel.

**Supplemental Figure 2.**
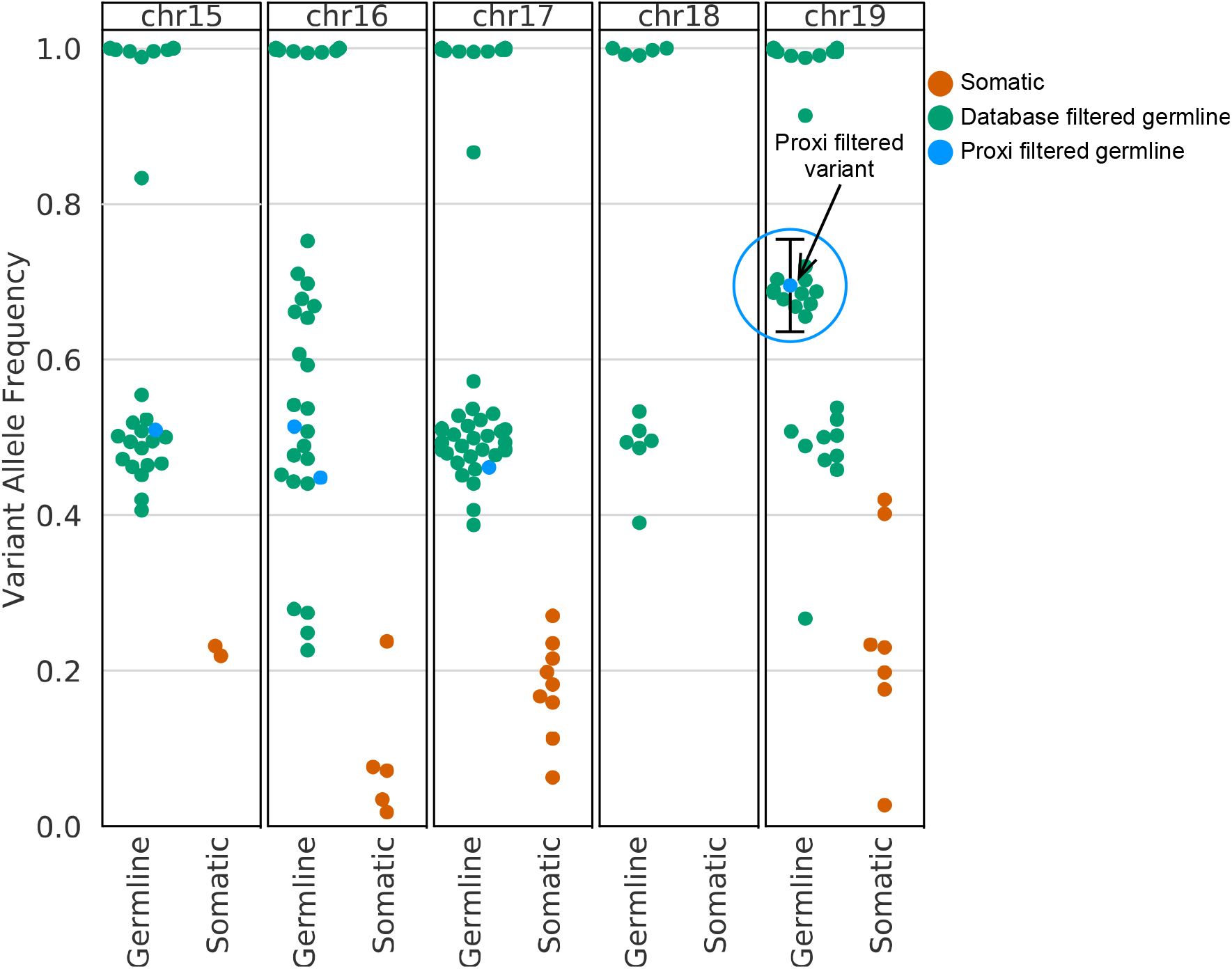
Germline variant filtering. Variants that are observed 10 times or more in public germline variant databases are filtered. Additional variants are categorized as germline variants using the proximity filter, which uses the database information of variants with a similar allele frequency (circled variants). Black bars indicate allele frequency range of the proximity filter, which is the maximum of 0.05 or 2 standard deviations giving a binomial distribution.

**Supplemental Figure 3.**
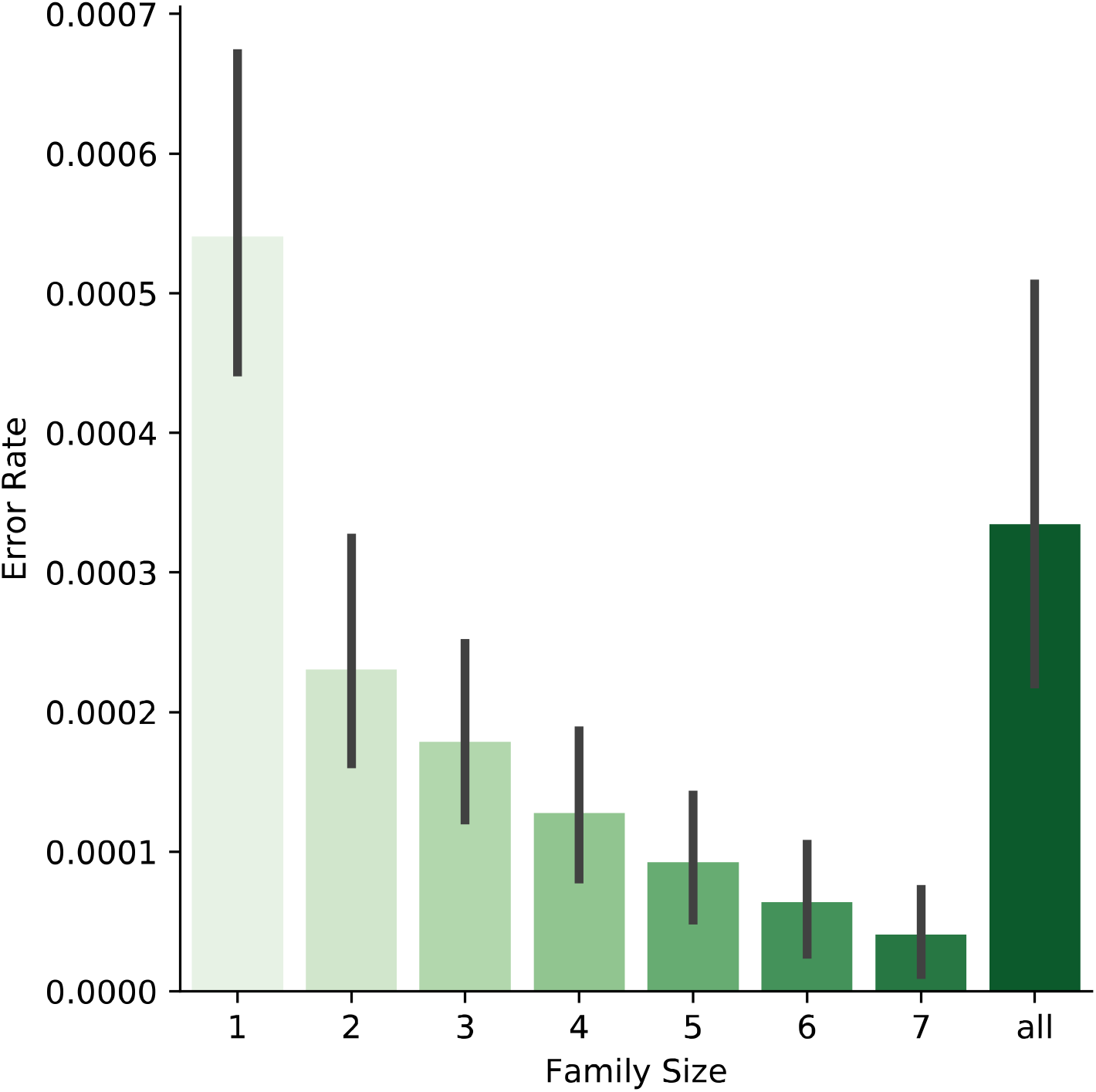
Sequencing error rate by family size. Read family size is indicated on the horizontal axis. The vertical axis gives the mean sequencing error rate for reads with a given family size. Black bars indicate the 95% confidence interval.

**Supplemental Figure 4.**
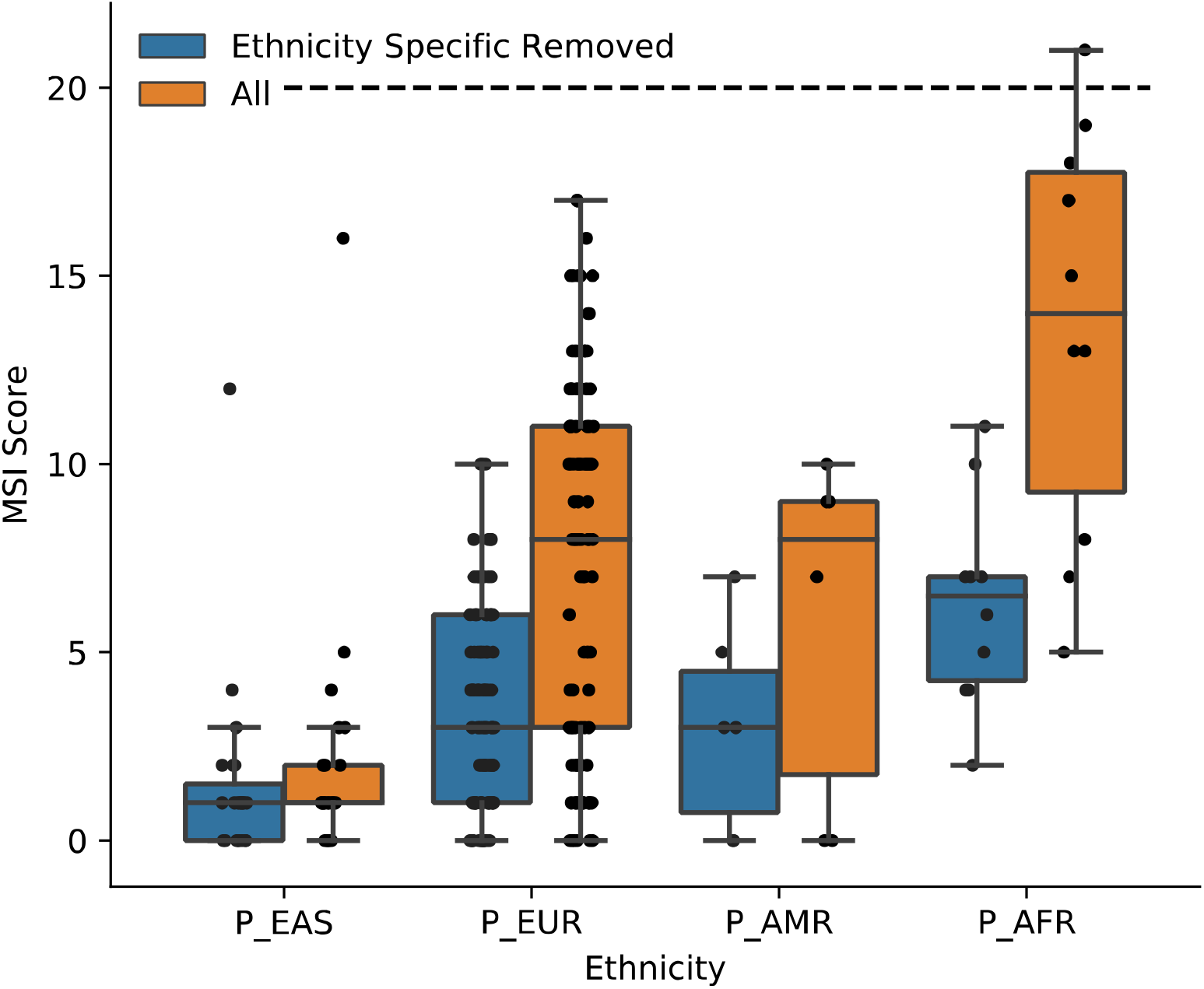
MSI scores for normal FFPE samples stratified by ethnicity. The horizontal axis gives the ethnicity of 140 normal FFPE samples assayed using TSO500. The vertical axis gives the MSI score calculated by the TSO500 bioinformatics workflow. The dashed line indicates the MSI score cutoff for determining MSI-high samples.

**Supplemental Figure 5.**
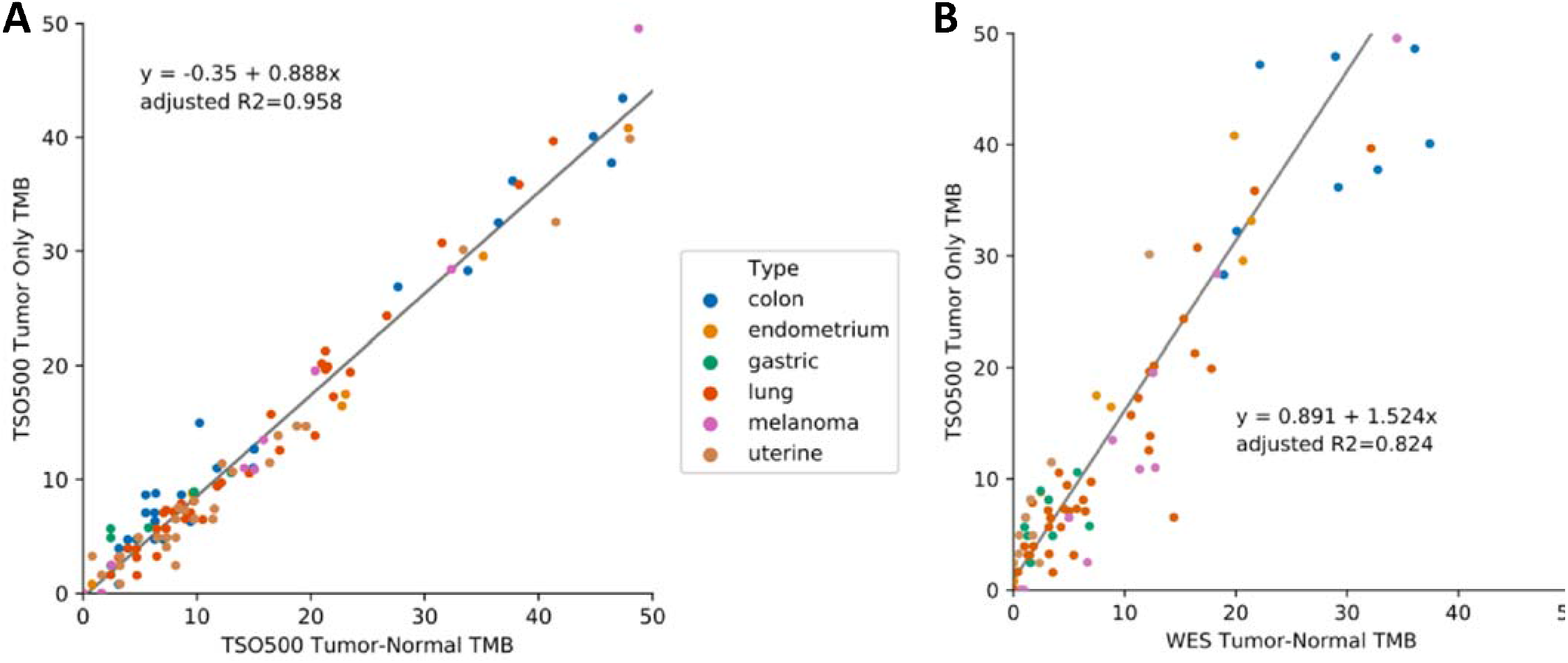
Comparison of TSO500 tumor-only TMB values to tumor-normal TMB values (zoomed in to TMB < 50).

**Supplemental Figure 6.**
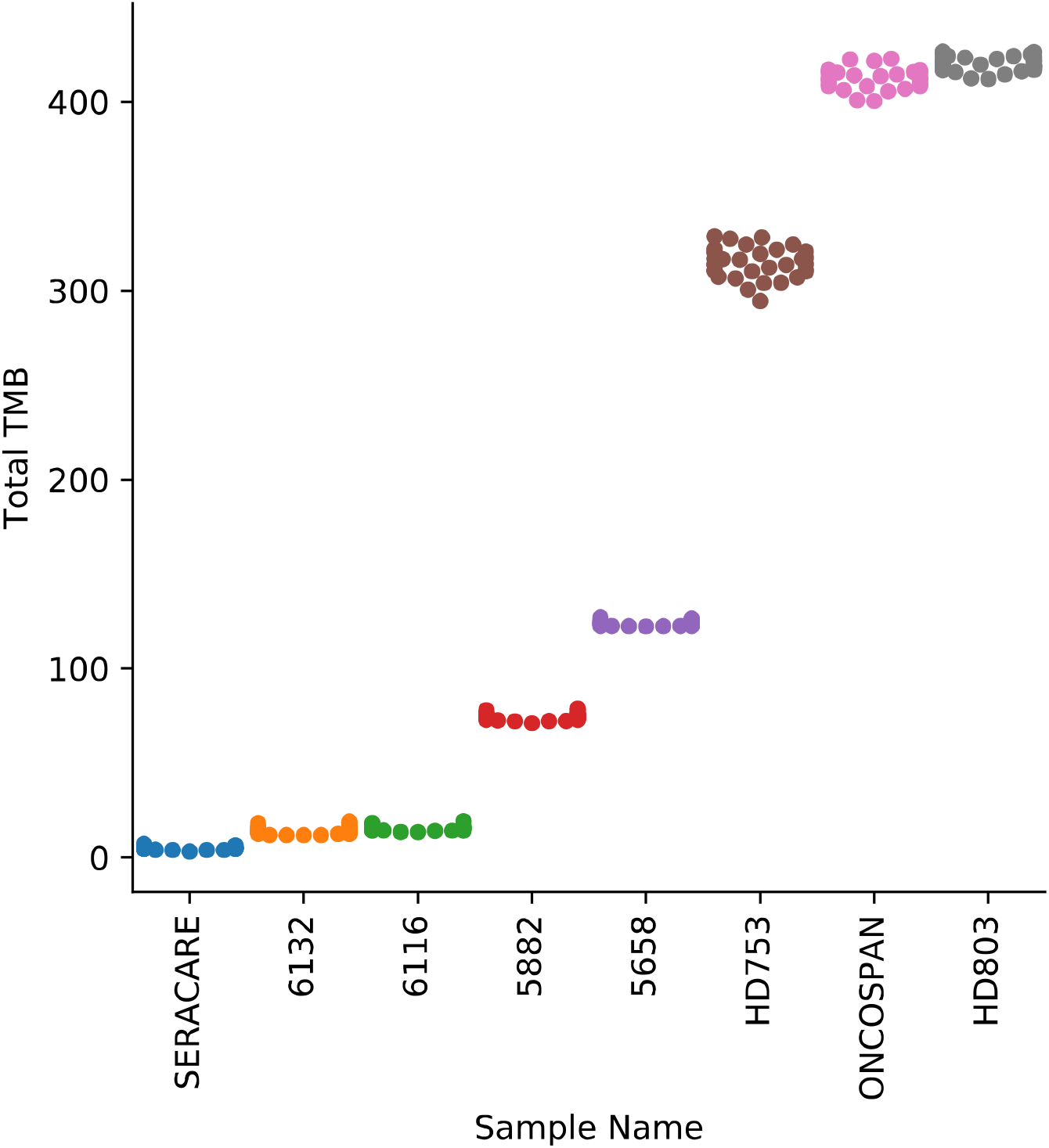
Reproducibility of TSO500 tumor-only TMB scores. Tumor-only TMB scores for 36 technical replicates of FFPE lung (6116, 6132), FFPE colon (5882, 5658), and commercial control (HD803, Oncospan, HD753, Seracare) samples are shown.

**Supplemental Figure 7.**
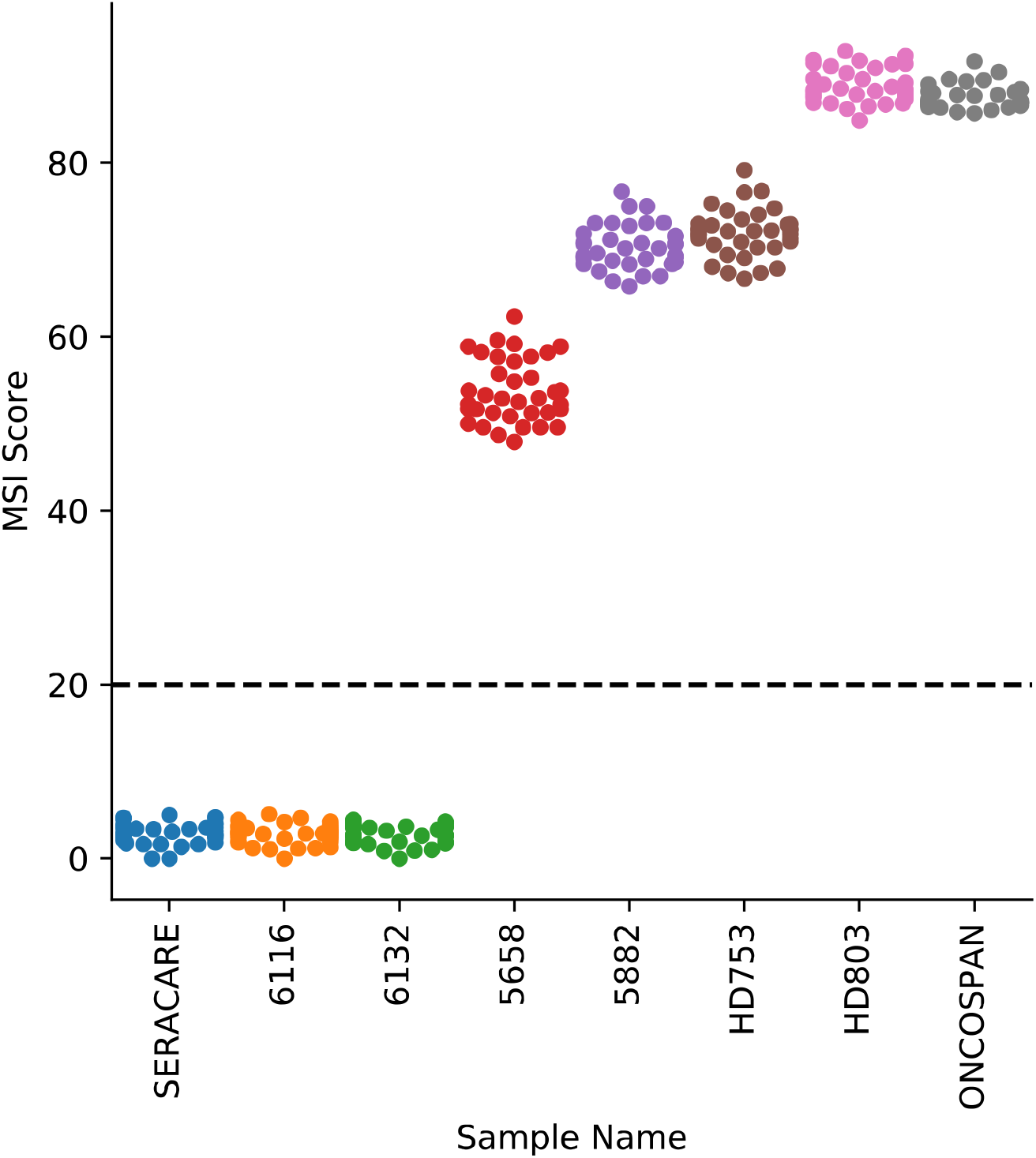
Reproducibility of TSO500 tumor-only MSI scores. Tumor-only MSI scores for 36 technical replicates of FFPE lung (6116, 6132), FFPE colon (5882, 5658), and commercial control (HD803, Oncospan, HD753, Seracare) samples are shown. The dashed line indicates the MSI score cutoff for determining MSI-high samples.

## Notes

### Competing Interest Statement

All authors are employees and shareholders of Illumina Inc.

