## Supplementary figures and images for "TruSight Oncology 500: Enabling Comprehensive Genomic Profiling and Biomarker Reporting with Targeted Sequencing"

### supplemental_figure_01

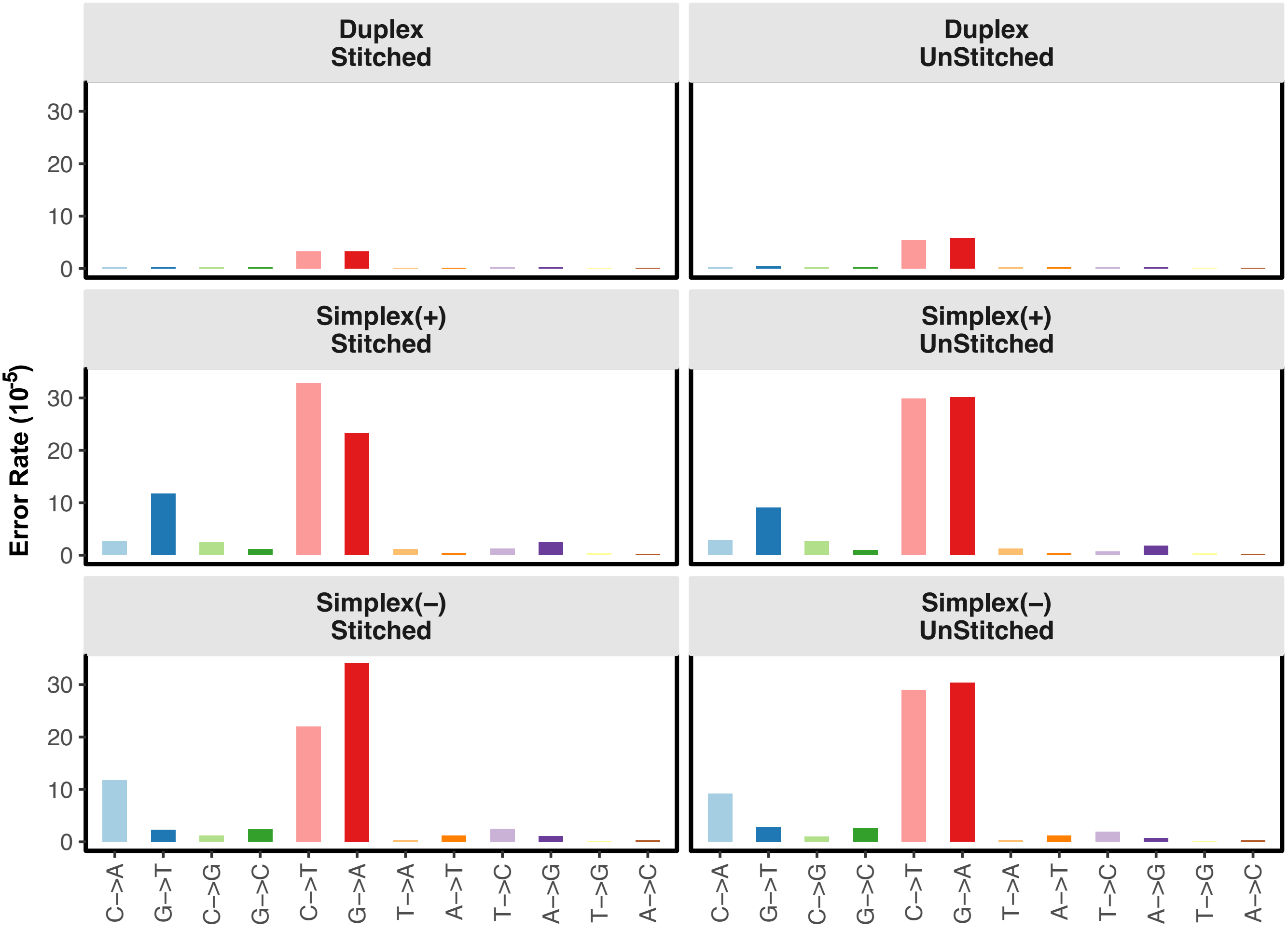

### supplemental_figure_02

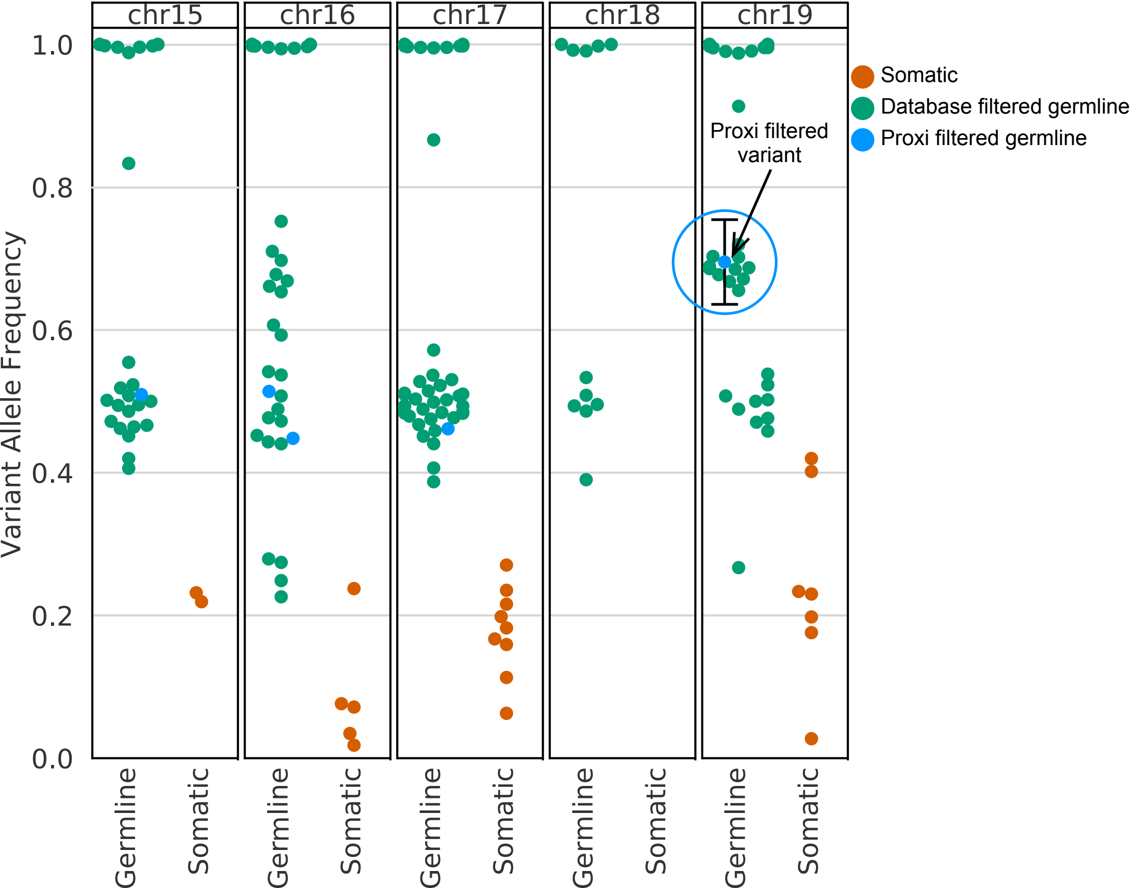

### supplemental_figure_03

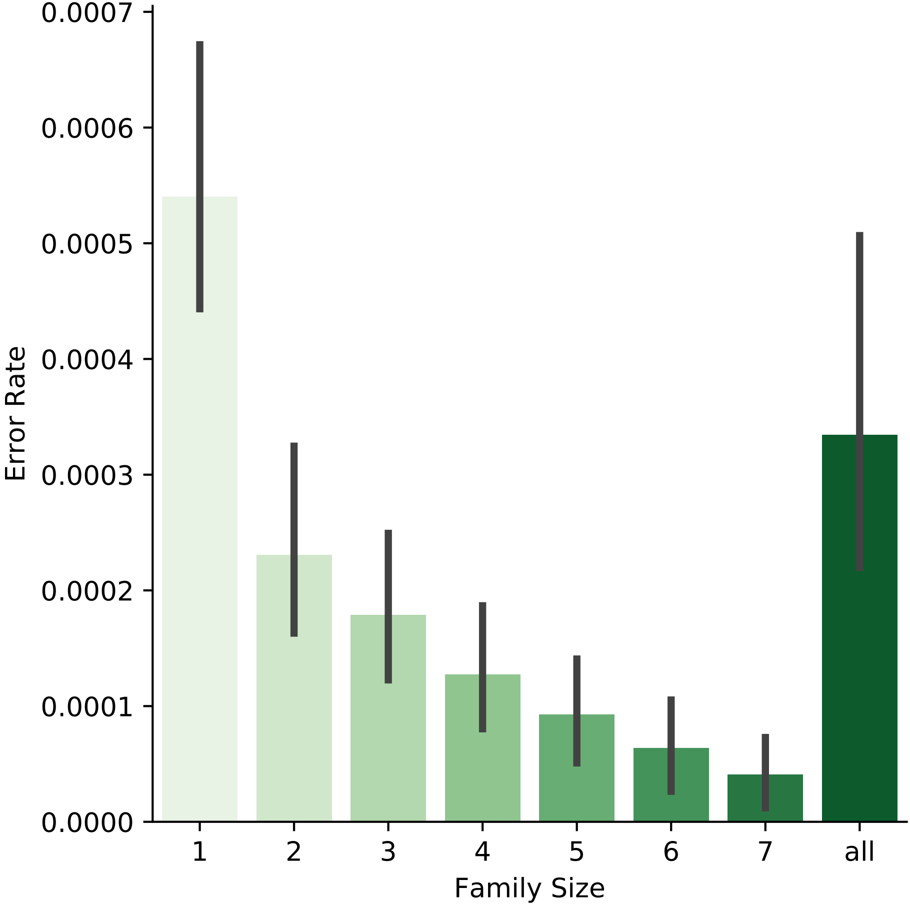

### supplemental_figure_04

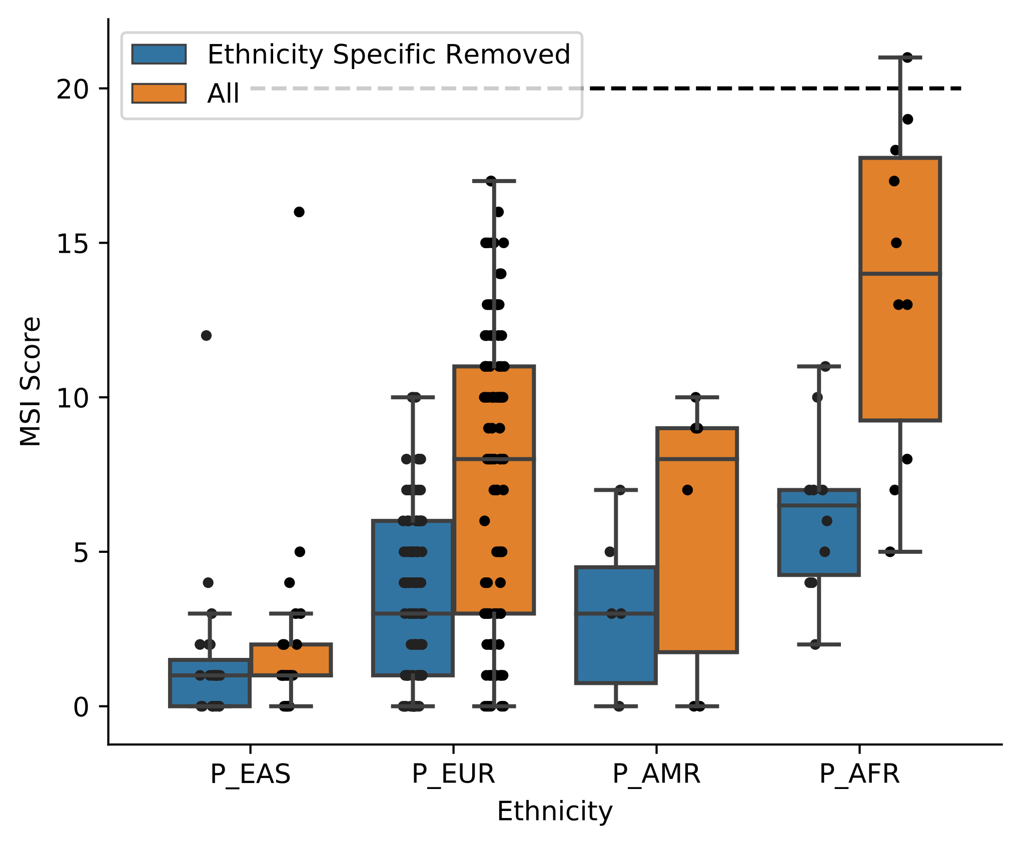

### supplemental_figure_05

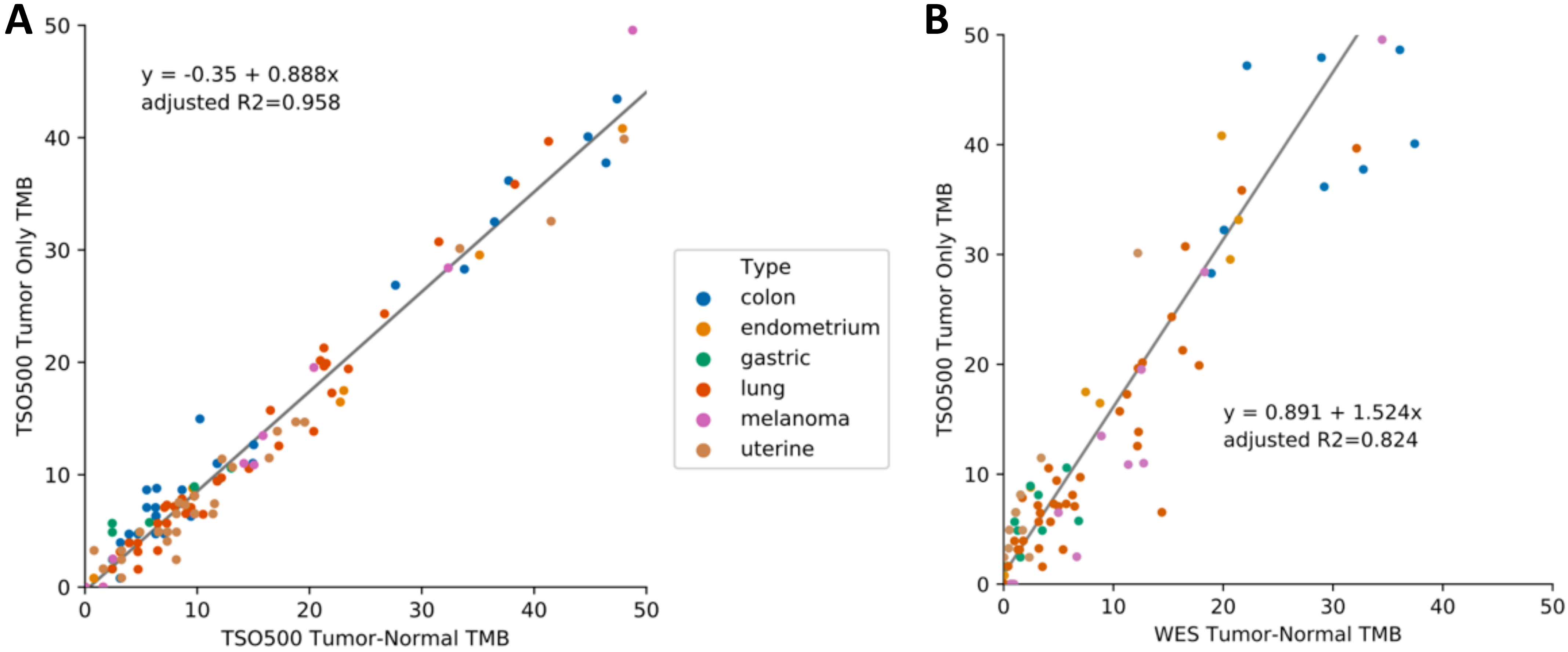

### supplemental_figure_06

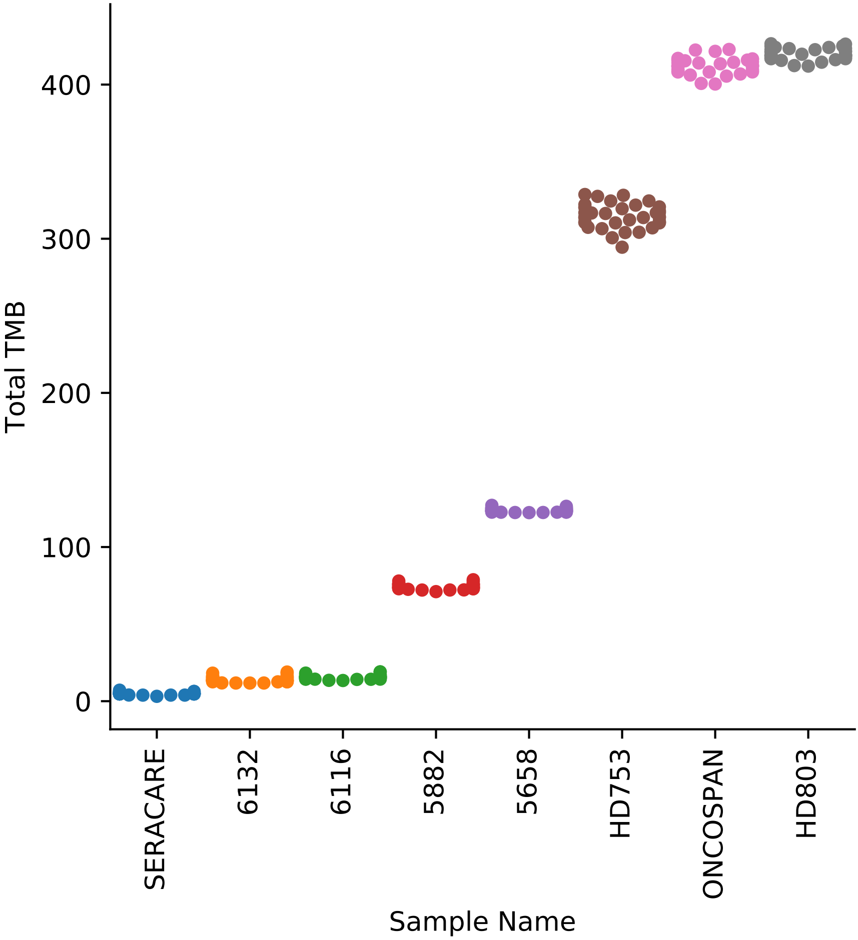

### supplemental_figure_07

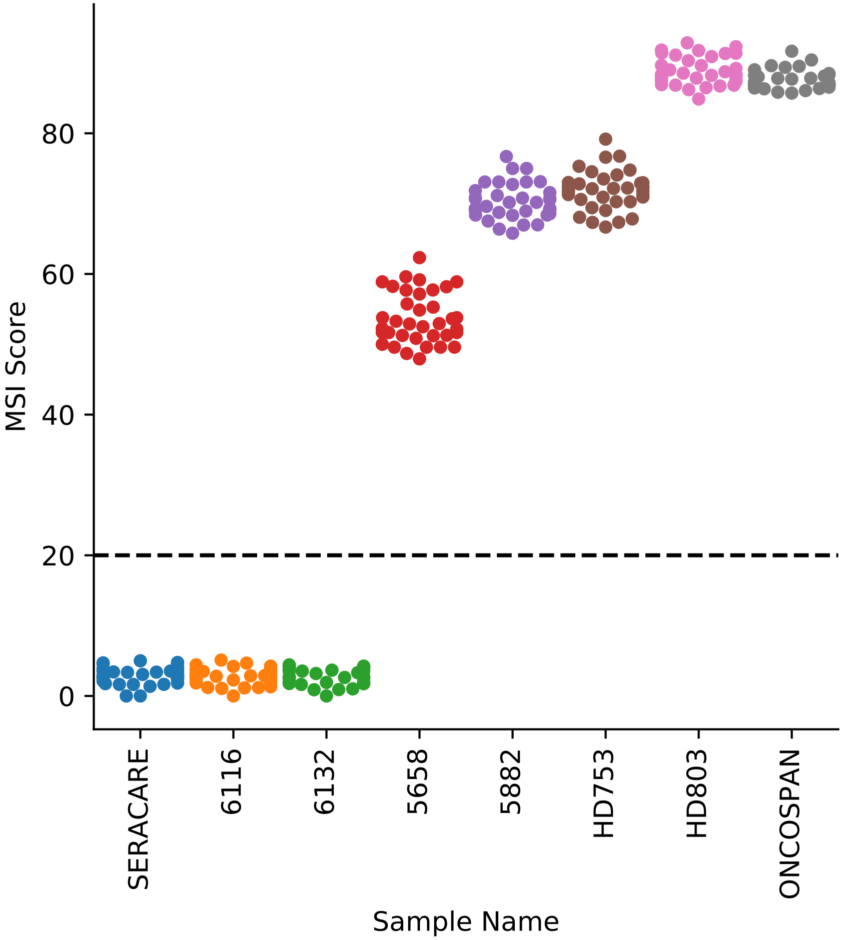
